# The desensitization pathway of GABA_A_ receptors, one subunit at a time

**DOI:** 10.1101/2020.05.18.101287

**Authors:** Marc Gielen, Nathalie Barilone, Pierre-Jean Corringer

## Abstract

GABA_A_ receptors mediate most inhibitory synaptic transmission in the brain of vertebrates. Following GABA binding and fast activation, these receptors undergo a slower desensitization, whose conformational pathway remains largely elusive. To explore the mechanism of desensitization, we used concatemeric α1β2γ2 GABA_A_ receptors to selectively introduce gain-of-desensitization mutations one subunit at a time. A library of twenty-six mutant combinations was generated and their bi-exponential macroscopic desensitization rates measured. Introducing mutations at the different subunits shows a strongly asymmetric pattern with a key contribution of the γ2 subunit, and combining mutations results in marked synergistic effects indicating a non-concerted mechanism. Kinetic modelling indeed suggests a pathway where subunits move independently, the desensitization of two subunits being required to occlude the pore. Our work thus hints towards a very diverse and labile conformational landscape during desensitization, with potential implications in physiology and pharmacology.

## INTRODUCTION

GABA_A_ receptors (GABA_A_Rs) are the main inhibitory synaptic receptors in the forebrain of vertebrates, and are involved in key physiological and pathological processes such as memory, epilepsy, anxiety, sedation. This is well-illustrated by their medical significance, since the most prevalent GABA_A_Rs are the target of the widely used benzodiazepine class of drugs^1^.

GABA_A_Rs belong to the pentameric Ligand-Gated Ion Channel (pLGIC) superfamily, which also comprises the anionic glycine receptor, as well as the excitatory 5HT_3_ serotonin receptors and the nicotinic acetylcholine receptors (nAChRs)^2^. Upon binding of their agonist, the transmembrane pore of pLGICs quickly opens to enable the selective flow of permeant ions across the plasma membrane, thereby affecting cell excitability. However, during sustained binding of the agonist, most pLGICs will gradually enter a shut-state refractory to activation, called the desensitized state, thereby preventing excessive activation^3^. The exact roles of desensitization *in vivo* are still debated, but potentially include the reduction of responses during high-frequency release of neurotransmitter^4^, the prolongation of synaptic currents due to a contribution of the recovery from desensitization^5^, as well as the modulation of extra-synaptic receptors subjected to tonic activation by low ambient concentrations of neurotransmitters^6^.

Recent functional and structural studies, mostly performed on anionic pLGICs, provide compelling evidence for a “dual-gate” model, in which the transmembrane domain (TMD) of pLGICs contains both an activation gate, located in the upper half of the channel, and a desensitization gate, located at the intracellular end of the channel^3,7–11^. Structural work on homopentameric receptors always showed symmetrical structures^7,9,11^, while the recent structures of the heteromeric GABA_A_ receptor show important asymmetric features within the extracellular domain (ECD)^10^, but still a strong pseudo-symmetrical organization of the TMD. The current view of the dual-gate model thus supports that resting, active and desensitized states are essentially symmetrical at the level of the TMD, desensitization involving, in the lower part of the channel, a movement of all subunits to occlude the permeation pathway. However, desensitization is a multiphasic process, since the sustained application of agonist elicits currents that desensitize with several distinct decay time constants, which are usually portrayed by the existence of “fast” and “slow” desensitized states (noted D_fast_ and D_slow_ below, respectively)^3,12–15^. The structural rearrangements underlying these distinct desensitization components remain elusive. In particular, it is currently unknown whether subunits rearrange in a concerted manner, with D_fast_ and D_slow_ reflecting distinct states at the single-subunit level, or whether individual subunits can rearrange independently with distinct time courses. The first scheme would predict that pLGICs only visit pseudo-symmetrical states during desensitization, while the latter scheme would imply that desensitization involves asymmetrical states.

To examine the contribution of individual subunits to this process, we herein introduced gain-of-desensitization mutations in each individual subunit, both one by one and in combinations, and assessed their interplay during desensitization. We selected mutations nearby the desensitization gate, which were previously found to specifically alter the desensitization kinetics and amplitude, without significant alteration of the upstream activation process. Since the stereotypical synaptic GABA_A_Rs are composed of two α, two β and one γ subunits^16,17^, targeting a single α or β subunit within the pentamer is out of reach using classical site-directed mutagenesis approaches. To circumvent this problem, we used a concatemeric construct, whereby all five subunits are linked together by polyglutamine linkers. Owing to the fixed organization of subunits within this concatemer, we could introduce and combine gain-of-desensitization mutations in a defined manner, ensuring the perfect homogeneity of the resulting recombinant GABA_A_Rs populations. We generated a library of 26 combinations of mutated subunits, recorded their macroscopic desensitization kinetics, and analyzed the data by Markov-chain kinetics simulations.

## RESULTS

### A pentameric concatemer recapitulates the biphasic desensitization profile of the GABA_A_R reconstituted from loose subunits

To force the subunit arrangement, we used a concatemer previously described^18^. It consists of β2-α1-β2-α1-γ2 subunits fused together with polyglutamine linkers that are 15- to 20-residues long. When assembled in the counter-clockwise orientation as seen from the extracellular space, it shows a canonical organization with two GABA binding sites at the β2-α1 interfaces and one benzodiazepine site at the α1-γ2 interface (Figure 1A). In contrast, in the clockwise orientation, the concatemer would carry a single GABA binding site and no benzodiazepine binding site. This orientation, if it occurs, should therefore yield minimal, if any, GABA-gated currents and no potentiation by benzodiazepine. We previously showed that expression of the concatemer in oocytes yields robust GABA-elicited currents and a potentiation by benzodiazepine indistinguishable from that of GABA_A_ receptors expressed from loose subunits^18^. This shows that the counter-clockwise assembly largely dominates the electrophysiological response.

**Figure 1.**
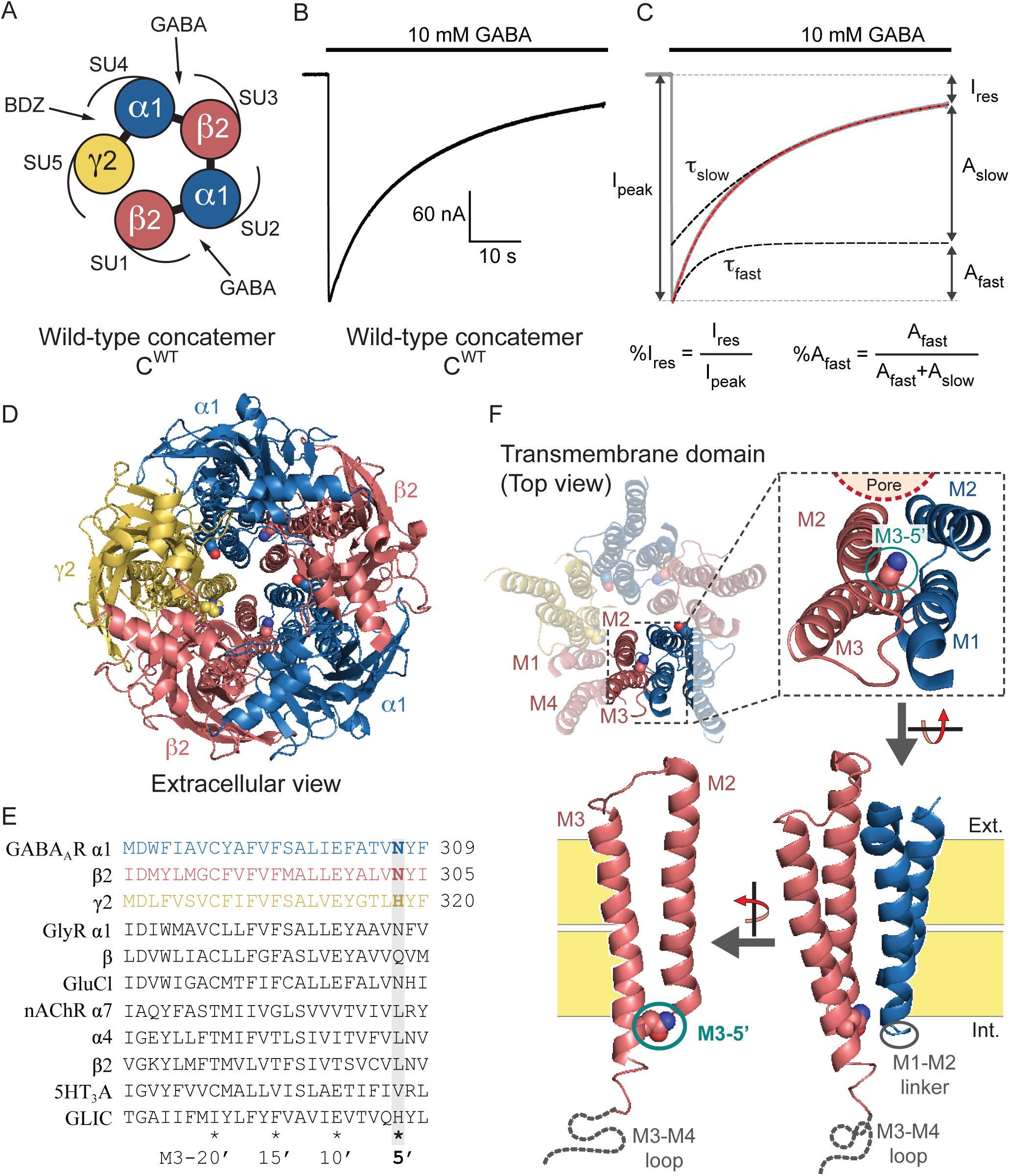
The wild-type α1β2γ2 pentameric GABA_A_R concatemer. ***A***, Schematic top view of the concatemer. The two β2/α1 ECD interfaces (SU1/SU2 and SU3/SU4) harbour the two GABA-binding sites, while the α1/γ2 ECD interface (SU4/SU5) contains the benzodiazepine-binding site. ***B***, Representative TEVC recording of a *Xenopus laevis* oocyte expressing the wild-type concatemer, C^WT^. ***C***, Depiction of the experimental values used to quantify desensitization: τ_fast_ and τ_slow_ are the time constants of fast and slow desensitization components, respectively; %A_fast_ is the relative amplitude of the fast component; %I_res_ is the relative residual current after 1 min of 10 mM GABA application. Of note, the weighted desensitization time constant can be defined as τ_w_ = %A_fast_ * τ_fast_ + (1-%A_fast_) * τ_slow_. ***D***, Cryo-EM structure of the α1β3γ2 GABA_A_R (pdb 6I53^17^), as seen from the extracellular space. The β2 and β3 GABA_A_ subunits are highly homologous, and both display an asparagine residue at the M3-5’ position. Note the central pore, lined by the M2 helices of the five subunits, forming the transmembrane channel. ***E***, Sequence alignment of the M3 segment of various pLGIC subunits. All sequences are the mouse orthologs, except GLIC (Gloeobacter violaceus), as well as the α4 and β2 nAChR subunits (human). The M3-5’ residues, mutated in the present study, are highlighted (grey box; bold characters for GABA_A_ subunits). ***F***, Enlarged view of the α1β3γ2 GABA_A_R structure highlighting the location of the M3-5’ residue at the M2/M3 transmembrane interface as seen from the side of the channel, facing the M1-M2 linker of the adjacent subunit.

To record desensitization kinetics at the best possible temporal resolution using Two-Electrode Voltage Clamp (TEVC) recordings of *Xenopus laevis* oocytes, we minimized the dead volume of our set-up and applied GABA at a supersaturating concentration (10 mM). This boosted the rise of GABA concentration in the recording chamber, enabling us to optimize the onset of the electrophysiological responses in the 20-25 ms timescale (corresponding to 20%-80% current rise times). Recordings of the wild-type concatemer show robust currents, which display desensitization kinetics and amplitudes similar to that of the conventional α1β2γ2 GABA_A_Rs assembled from unconnected subunits (Figure 1B&C; Supplementary Table 1; see ref^8^). Desensitization shows two well-separated components that are perfectly resolved by our procedure, a fast (τ_fast_ = 4.8 ± 1.2 s) and a slow one (τ_slow_ = 24.4 ± 7.8 s). The amplitude of the former carries about a third of the total desensitization amplitude, both fast and slow rates together yielded a weighted desensitization time constant (τ_w_) of about 18 s. The residual current remaining after one minute of GABA application accounted for about 10% of the peak current (Figure 1B&C; Supplementary Table 1).

### Single desensitizing mutations show contrasting phenotypes depending on their location within the pentamer

For gain-of-desensitization mutations in α1, β2 and γ2 subunits, we chose the valine mutation at the 5’ position of the third transmembrane segment (M3; Ref^19^), namely α1^N307V^ on α1 subunits (SU2 and SU4), β2^N303V^ on β2 subunits (SU1 and SU3) and γ2^H318V^ on the single γ2 subunit (SU5) (Figure 1D-F). Indeed, on GABA_A_Rs composed of loose subunits, we previously showed that these mutations markedly speed up the onset of desensitization of α1β2γ2 GABA_A_Rs^8^. We also showed that mutations in this region of the transmembrane domain do not alter significantly the concentration-response curve of the GABA-elicited peak currents measured before the onset of desensitization. This indicates only a weak effect of the mutations on the resting-to-active state transition, and a major effect on the active-to-desensitized state transition.

Mutations were introduced one at a time on the concatemer. We define C^WT^ as the wild-type concatemer, C^*i*^ the concatemer with a single M3-5’ valine mutation on subunit number *i*, and C^*ij*^ the concatemer where subunits *i* and *j* are both mutated, up to C^12345^ where all subunits are mutated (Figure 2A).

**Figure 2.**
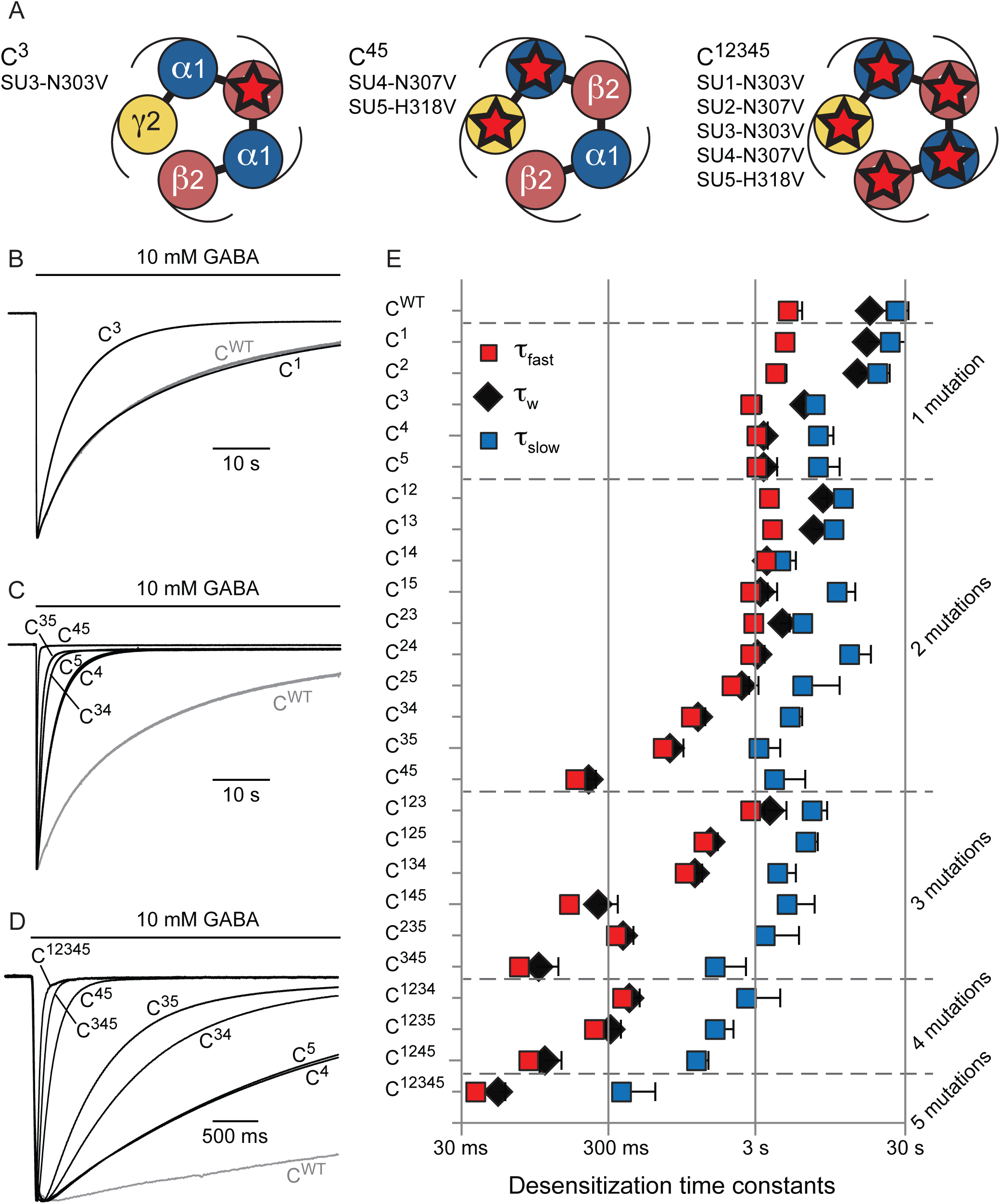
Desensitization kinetics of α1β2γ2 concatemers harbouring combinations of M3-5’ valine mutations. ***A***, Schematic top views of the C^3^ (left), C^45^ (middle) and C^12345^ (right) concatemers. ***B-D***, Representative TEVC recording of *Xenopus laevis* oocytes expressing the indicated concatemers. Note the change in timescale for recordings in panel ***D. E***, Plot indicating the fast (red squares), slow (blue squares) and weighted (black diamonds) desensitization time constants for the indicated concatemers. Error bars are standard deviations.

For the single mutations, C^1^ (SU1 = β2) and C^2^ (SU2 = α1) display desensitization kinetics similar to that of C^WT^, while constructs C^3^, C^4^ and C^5^ displayed robust gain-of-desensitization phenotypes (Figure 2; Supplementary Table 1), yielding weighted desensitization rates of 6.2, 3.4 and 3.3 s, respectively, as compared to 18 s for C^WT^. The three mutations accelerate fast desensitization by about 2-fold and slow desensitization by about 3-fold (τ_fast_ = 2.7, 2.9 and 2.9 s; τ_slow_ = 7.1, 7.3 and 7.2 s for C^3^, C^4^ and C^5^, respectively). C^4^ and C^5^ in addition increase the relative amplitude of the fast component (%A_fast_ = 20.0%, 86.7% and 86.3% for C^3^, C^4^ and C^5^, respectively), explaining their stronger effect.

It is noteworthy that the mutations are located at the cytoplasmic end of the TMD, with the side-chain of the mutated residue facing the M1-M2 linker of the neighbouring subunit (Figure 1F). Therefore, C^1^, C^2^, C^3^, C^4^ and C^5^ are mutated at β2-α1, α1-β2, β2-α1, α1-γ2 and γ2-β2 interfaces, respectively. The different mutations being introduced at different interfaces, it was expected that they display different phenotypes. However, the difference between C^1^ and C^3^ is surprising, since they both correspond to mutations at the β2-α1 interface, showing virtually identical microenvironment. This indicates that the effect of the single mutations not only depends on the nature of the mutated interface, but also on the particular position of the mutated subunit within the pentamer.

### Combining mutated subunits increases desensitization kinetics and reveals synergistic effects

To investigate the functional interaction between mutations at the various interfaces, we built an extensive library of twenty-six cDNAs including concatemers comprising two mutations (ten different constructs), three mutations (six different constructs), four mutations (four different constructs) or five mutations (one single construct, C^12345^), and assessed their desensitization profile as described above (Figure 2; Supplementary Table 1).

Recordings confirmed the modest effect of SU1 and SU2 mutations, which produce small effects when performed on concatemers with background mutations at other subunits (0.8 to 1.8-fold decrease in τ_w_ for SU1 and 1.1 to 2.8-fold for SU2, among 9 background mutated concatemers for both). They also confirm the intermediate effect of SU3 (2 to 6.3-fold decrease in τ_w_ among 10 background mutated concatemers), and the marked effect of SU4 and SU5 (effect of 5 to 16-fold among 9 and 10 mutated concatemers for SU4 and SU5, respectively; Supplementary Figure 1).

In all cases, combining gain-of-desensitization mutations together adds up to increase desensitization kinetics. For instance, the double mutant C^45^ displays a fast desensitization component (τ_fast_ = 180 ms) 26-fold faster than C^WT^, accounting almost entirely for the overall desensitization (%A_fast_ = 98.8%), and a barely measurable steady-state current (%I_res_ = 0.8%). Such phenotype is further strengthened by mutating SU3: C^345^ desensitizes with an even faster desensitization component in the 70 ms timescale. Mutating all five subunits gave a slightly more profound phenotype, with a fast desensitization component of 40 ms (see construct C^12345^; Figure 2D & E; Supplementary Table 1). Of note, for constructs akin C^345^ and C^12345^, the fast component is so fast that we probably miss a sizeable fraction of the peak current, thereby overestimating the amplitude of the slow desensitization component and the measurement of the relative steady-state current. Also, the steady-state current values and the amplitudes of the slow desensitization components are barely measurable for constructs displaying the fastest and most complete desensitization, rendering the related values (%Ires and τ_slow_) unreliable.

To investigate the additivity of the effect of the various mutations, we first compared the effect of individual mutations on the weighted desensitization kinetics of different concatemers with background mutations (Supplementary Figure 1). While this analysis is crude, the series of double mutants already suggests some level of inter-subunit coupling. Indeed, while the SU1 mutation barely affects the desensitization of C^WT^, it increases the weighted desensitization kinetics of C^2^ by 75%, thereby hinting towards a coupling between SU1 and SU2. More strikingly, SU4 mutation speeds up desensitization about 5-fold on both C^WT^, C^1^, C^2^ and C^3^ backgrounds, while it increases the weighted desensitization kinetics of C^5^ by 15-fold, clearly hinting towards synergistic effects of SU4 and SU5 mutations.

Second, we compared the desensitization profiles of C^34^ and C^35^. Since mutating SU4 or SU5 yields identical desensitization phenotypes (Figure 2C-E; Supplementary Table 1), C^34^ and C^35^ should yield identical phenotypes if the effects of mutations were additive. Our data contradict such hypothesis, since both desensitization components of C^35^ are faster than the ones of C^34^, resulting in a 55% faster weighted desensitization rate (Figure 2C-E; Supplementary Table 1). Thus, the effects of mutating the M3-5’ residues are non-additive, especially for SU3 and SU4 or SU3 and SU5 subunit combinations.

### The conformational pathway of desensitization involves asymmetrical and non-concerted quaternary motions: implementation of a general model

The present analysis unravels two key features governing the desensitization kinetics. First, the markedly different effects observed upon mutation of SU1 and SU3, which both involve homologous mutations that are located in identical micro-environments, show that strongly asymmetrical motions are involved in the desensitization pathway. Since SU3 mutation has a strong effect on desensitization, the structural reorganization at this interface appears to be a limiting process. In contrast, mutation in SU1 has very weak effect, suggesting either a small structural reorganization at this level, or, more likely, that the structural reorganization would not be rate limiting (see Discussion).

Second, the marked non-additive nature of the mutations, as discussed above, is not compatible with a concerted mechanism. Indeed, in such a scheme, the effect of mutations should directly translate their impact on the free energy landscape of the receptor, and should thus be additive.

As an illustration, we attempted to fit the whole set of data with a concerted model, in which the receptors can only visit a handful of pseudo-symmetrical conformations that include a fast and a slow desensitized state (Supplementary Figure 2A-E). Here and throughout the manuscript, each model was built as a Markov-chain kinetic scheme and the whole-cell currents activated by a supersaturating concentration of GABA were simulated using the software QUB^20^ (Supplementary Table 2). However, adjusting the parameters to correctly fit the desensitization of C^WT^, C^4^ and C^5^, did not account for their synergistic effect since the simulated C^45^ τ_fast_ and τ_w_ values are respectively 4.7 and 4.1-fold higher than the values observed experimentally (Supplementary Figure 2F-G).

To implement the asymmetric and non-concerted properties, we turned to a radically different scheme in which all subunits can desensitize independently from the other subunits (Figure 3). In this model, each subunit can enter its desensitized conformation while the other subunits are either in their open or desensitized conformations. For simplicity, we decided to implement only the desensitization of SU3, SU4 and SU5, since these subunits are by far the main contributors to the phenotypes in the dataset. This enabled us to reduce the model to ten different states, rather than thirty-four distinct states involving all subunits. We also strongly simplified the activation transition whereby the resting receptor (R state) binds the agonist (AR state) and subsequently open (AO state). The model thus does not account for unliganded receptors openings (O state) that rarely occur at wild-type α1β2γ2 GABA_A_Rs, with a spontaneous open probability as low as 10^−5^ in the absence of agonist^21^, nor does it include the binding of two GABA molecules: we only considered the gating equilibrium for fully occupied receptors, as we work with supersaturating concentrations of GABA. From the AO state, either SU3, SU4 or SU5 can desensitize, to produce AD_3_, AD_4_ or AD_5_ states, respectively. From these, the receptor can be further driven into states where two subunits are desensitized, e.g. desensitization of SU5 from the AD_4_ state leads to the AD_45_ state, where both SU4 and SU5 are desensitized. Finally, in that instance, SU3 could also desensitize to yield the AD_345_ state, in which all three subunits are desensitized.

**Figure 3.**
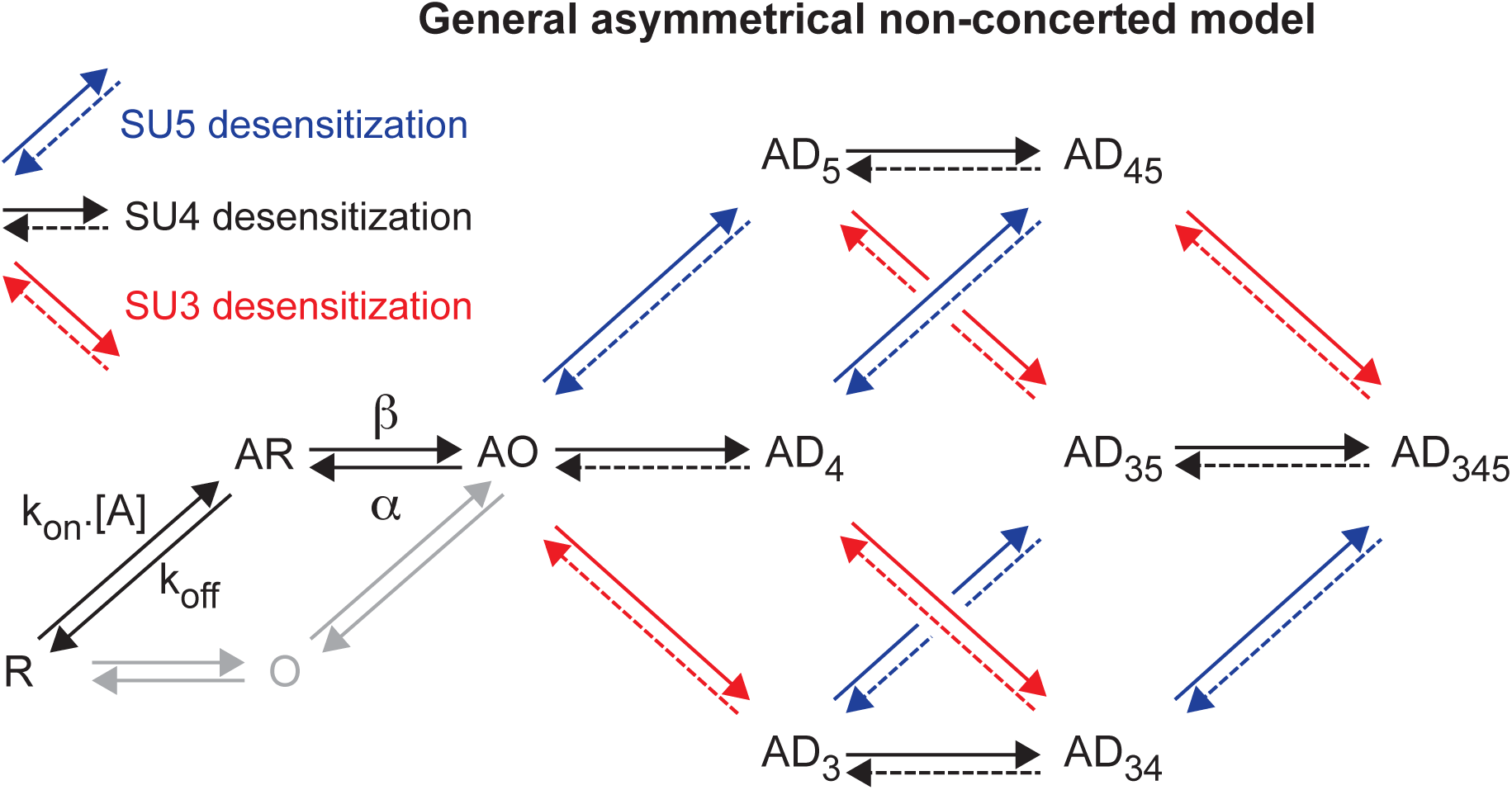
General scheme for the simulation of desensitization: an asymmetric non-concerted model. The first part in the kinetic scheme is the binding of the agonist A to the resting state R, which favours the opening of the channel (AO state) with a gating efficacy E = β/α. Of note, unliganded openings do exist but are not taken into account for our kinetic modelling as they barely contribute to the electrophysiological response (see main text). We also only include one binding event, even though α1β2γ2 GABA_A_Rs contain two binding sites whose occupation is required for substantial activation. Upon channel opening, the receptor can then transit from a fully activated AO state to states where only one subunit enters its desensitized conformation (AD_3_, AD_4_ and AD_5_). From these states, a second subunit can also desensitize, before the final step leading to the state in which all subunits are desensitized.

Using this general model, we progressively tuned the kinetic and functional parameters to best fit the dataset.

### Model I, in which desensitization of a single subunit shuts the channel, shows anti-synergistic behaviour

We first postulated that the receptor is functionally desensitized, i.e. non-conducting, as soon as one subunit is desensitized, with only the AO state allowing the passage of ions (Figure 4A).

**Figure 4.**
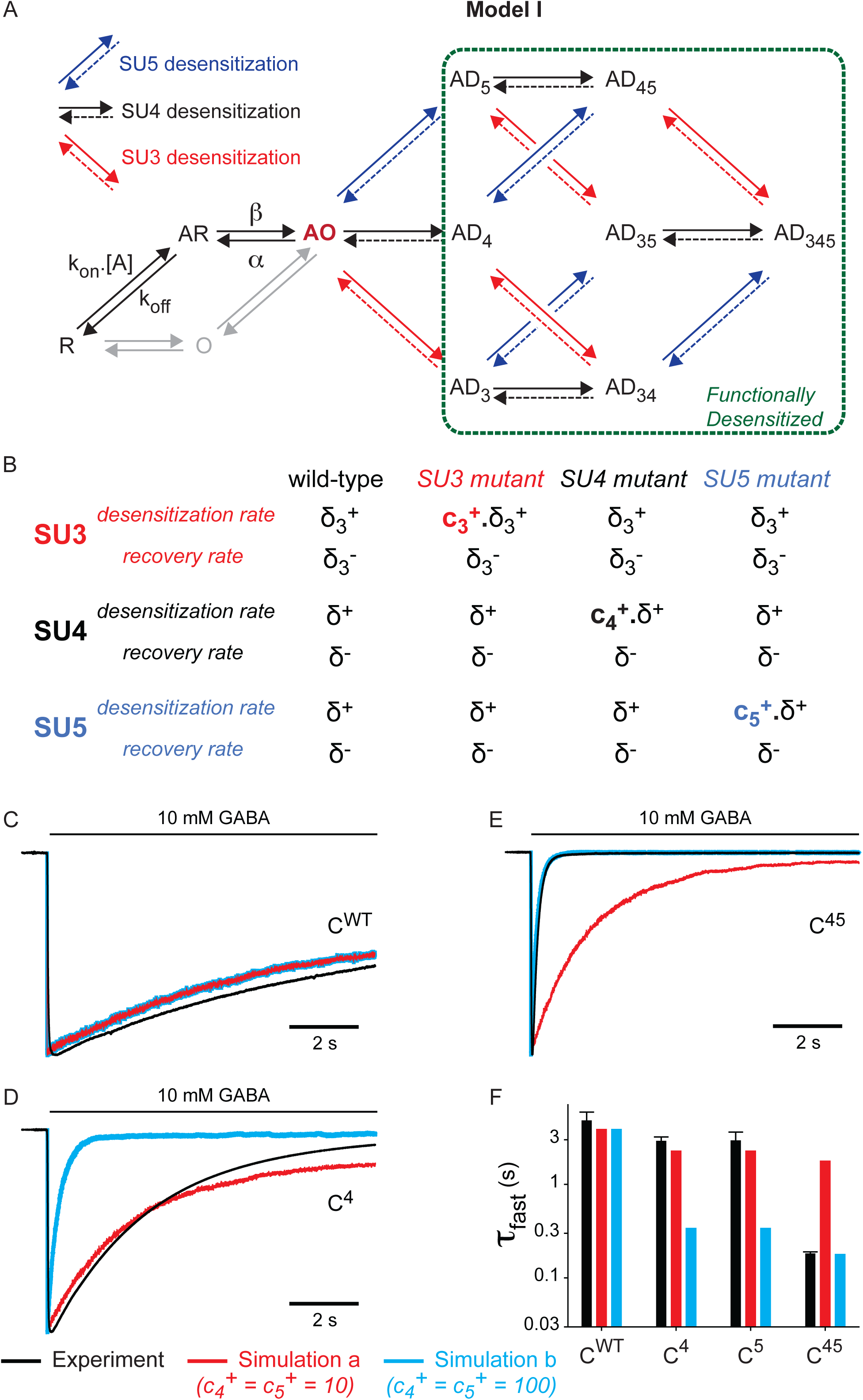
Model I: only the fully open state is conducting, and subunits move independently during desensitization. ***a***, We assume in this model that a single desensitized subunit is enough to shut the pore of the channel, leading to functional desensitization. Moreover, subunits SU3, SU4 and SU5 can undergo a desensitization rearrangement independent of the other subunits. Thus, desensitization rates (δ_3_^+^ for SU3, δ^+^ for SU4 and SU5) and recovery rates (δ_3_^-^ for SU3, δ^-^ for SU4 and SU5) do not depend on the conformation of the neighbouring subunits. ***B***, Effect of M3-5’ valine mutations in Model I. Mutations are hypothesized to specifically increase the desensitization rates of the mutated subunits, without altering any other parameter. ***C-E***, Representative currents for C^WT^ (panel ***C***), C^4^ (panel ***D***) and C^45^ (panel ***E***), in black, are compared to the outcome of two distinct simulations. In simulation *a* (red), the mutation-induced increase in the desensitization rates of SU4 and SU5 is adjusted so that the simulation of single mutants C^4^ and C^5^ broadly fits the experimental data, as seen in panel ***D***. In simulation *b* (blue), the mutation-induced increase in the desensitization rates of SU4 and SU5 is adjusted so that the simulation of the double mutant C^45^ accounts for the experimental data, as seen in panel ***E. F***, Bar graph summarizing the experimental data vs the predicted effects of SU4 and/or SU5 mutations on the kinetics of the fast desensitization component in simulations *a* and *b*. For panels ***C-F***, note that parameters from simulation *a* fail at describing the data for the double mutant C^45^, while parameters from simulation *b* largely overestimate the effect of single mutants. See Supplementary Table 3 for the numerical values of parameters.

In model I, the desensitization and recovery rates (δ^+^ and δ^-^) for each subunit do not depend on the state of the other subunits. For simplicity, the parameters for SU4 and SU5 are set equal, since C^4^ and C^5^ display similar phenotypes. Thus, only four parameters (δ^+^, δ^-^, δ_3_^+^and δ_3_^-^) are used to constrain the desensitization of C^WT^, i.e. exactly the number of independent numerical constraints provided by the experimental data (τ_fast_, τ_slow_, %A_fast_, %I_res_). We also assumed that mutating subunit *i* simply increases its desensitization rate by a ratio c_*i*_^+^ (Figure 4B).

For each set of parameters, we performed kinetic simulations using QUB (Figure 4 and Supplementary Table 3). Data are then analyzed using bi-exponential fitting of each virtual recording. In every simulation, we included all combinations of SU3, SU4 and SU5 mutants, from C^WT^ to C^345^.

In simulation “***a***”, we set up the parameters to reproduce C^WT^ and single mutant concatemers (Figure 4C, D & F and Supplementary Table 3). However, these parameters largely underestimate the kinetics of the fast desensitization component for the double mutant C^45^: simulation ***a*** predicts a value of 1.76 s for the τ_fast_ of C^45^, i.e. 10-fold slower than the experimental value. In simulation “***b***”, we used the same parameters for C^WT^, and set up the c_*i*_^+^ ratios to reproduce the C^45^ phenotype (Figure 4C, E & F and Supplementary Table 3). In that situation, we now largely overestimate the kinetics of the fast desensitization component for the single mutants C^4^ and C^5^: simulation ***b*** predicts a value of 0.34 s for the τ_fast_ of both C^4^ and C^5^, i.e. an order of magnitude faster than the experimental values. In this particular example, it is striking that model I actually predicts *anti-synergistic* effects when mutating SU4 and SU5, with the fast desensitization kinetics of both the single and double mutants being similar (Figure 4F).

Model I is thus incompatible with the dataset, and the reason is actually straightforward: if one desensitized subunit is enough to shut the pore, there should be a limiting fast subunit, whose mutation should have a strong effect on the kinetics of the fast desensitization component. This is not what we observe experimentally: the single mutant concatemers with the strongest phenotypes, C^4^ and C^5^, only display 40% increases in τ_fast_ (see above).

### Model II, in which at least two desensitized subunits are required to shut the pore, accounts for the synergy between SU4 and SU5 mutations

We consequently modified the kinetic model to incorporate a key hypothesis: namely, that functional desensitization of the channel involves the rearrangement of at least two subunits, i.e. that AO, AD_3_, AD_4_ and AD_5_ do conduct ions, while AD_34_, AD_45_, AD_35_ and AD_345_ do not (model II, Figure 5A).

**Figure 5.**
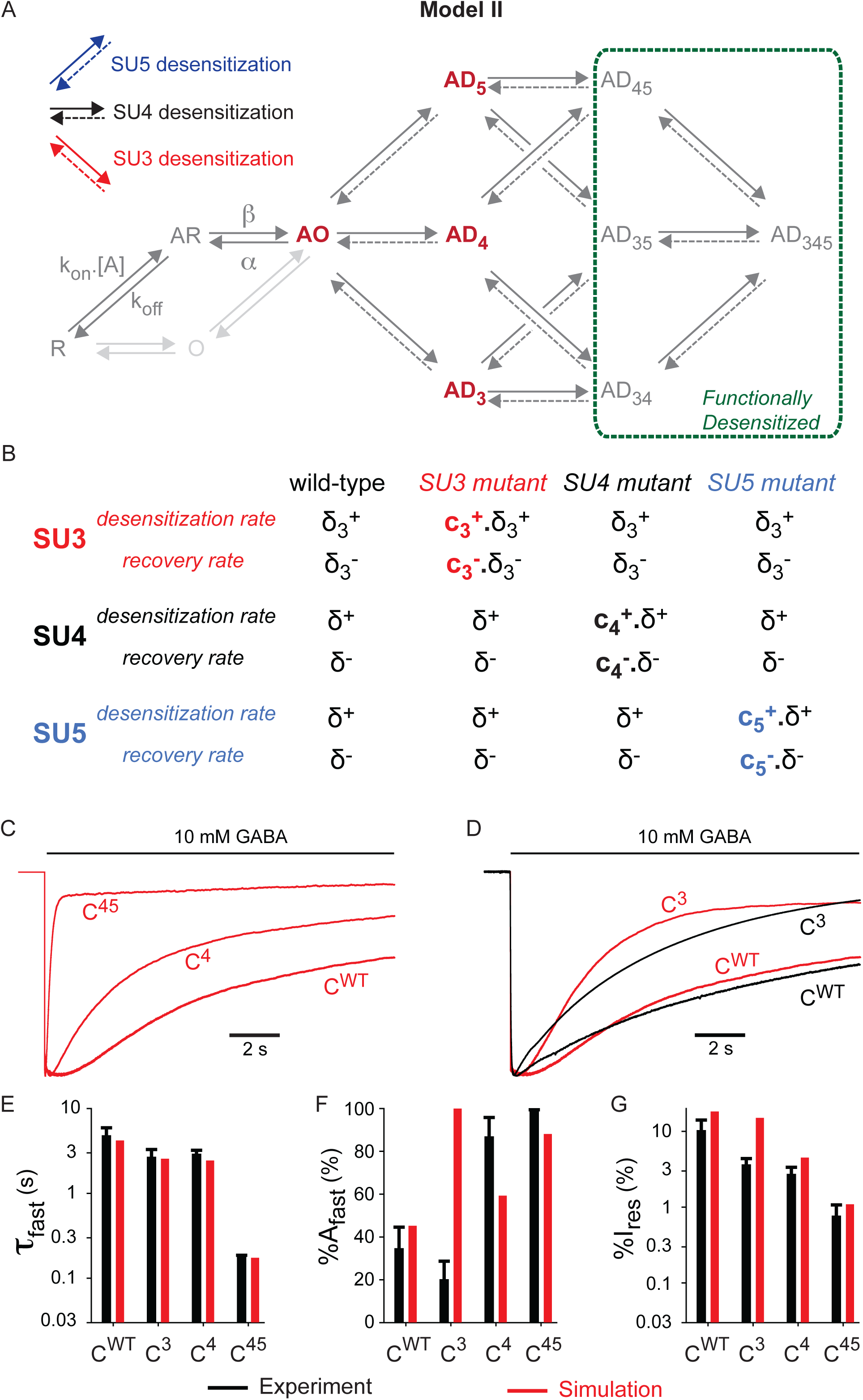
Model II: Two desensitized subunits are required to occlude the pore. ***A***, Model II builds upon Model I by adding one key hypothesis: receptors with only one subunit in its desensitized conformation are still conducting, and desensitization occurs when at least two subunits are desensitized. Thus, states AD_3_, AD_4_ and AD_5_ are open states from a functional point of view. ***B***, In Model II, mutation of a subunit can affect both its desensitization and recovery, as shown here with an example in which both SU4 and SU5 are mutated (construct C^45^): c_4_^+^ and c_5_^+^ reflect the increase in desensitization rates, c_4_^-^ and c_5_^-^ reflecting the increase in recovery rates. ***C***, Simulated currents for C^WT^, C^4^ and C^45^. ***D***, Representative currents for C^WT^ and C^3^ in black, are compared to their simulation counterparts in red. ***E-G***, Bar graphs summarizing the experimental data (in black) vs the simulations (in red) for the indicated concatemers on the kinetics (panel ***E***) and the amplitude (panel ***F***) of the fast desensitization component as well as the residual current after a 1 min long application of 10 mM GABA (panel ***G***). Note that the results for the C^5^ construct are not displayed, since the experimental data are almost identical to that of C^4^ (see Figure 2) and since the simulations for C^4^ and C^5^ are identical (see Supplementary Table 3). See Supplementary Figure 2 for all simulation results, and Supplementary Table 3 for the numerical values of parameters.

To simulate responses with steady-state currents consistent with experimental values, we also allowed mutations to increase the rates for desensitization recovery of the mutated subunits (Figure 5B; Supplementary Table 3). Using this model II, we could perfectly account for the fast desensitization rate of C^4^, C^5^ and C^45^ (Figure 5C-E). When SU4 is mutated, SU5 desensitization still provides a limiting step for functional desensitization, acting as a brake, while in C^45^ both “brakes” are relieved, enabling the channel to desensitize with fast kinetics, thereby generating a synergistic effect. This serves as a gentle reminder for studies using mutant-cycle analysis: it is indeed possible to have a strong functional coupling between non-interacting residues located far apart in a receptor’s structure, if their motions are not concerted.

While model II accounts for the main features of the dataset, we further refined it to precisely fit some desensitization kinetics. Indeed, simulation of C^3^ shows a mono-exponential process with %A_fast_ = 100% (Figure 5D & F), and an overestimated residual current (Figure 5G; Supplementary Figure 3). To circumvent this issue, we assumed that mutating SU3 increases the desensitization and recovery rates of SU4 (model II-β; Supplementary Figure 4). From a structural point of view, such hypothesis seems plausible: the M3-5’ residue mutated in SU3 is located at the interface with SU4 (Figure 1A&D-F), potentially interfering with conformational rearrangements of SU4. Using this model II-β, we could correctly simulate C^3^ with two components for desensitization, (Supplementary Figure 5A & B and Supplementary Table 3).

Still, for C^WT^ and C^3^, model II-β produces an overestimation of both the fast component amplitude and the residual current (Supplementary Figure 5C&D). Increasing the desensitization equilibrium constant (δ^+^/δ^-^) for SU4 and SU5 would reduce the amount of residual current, but would also lead to an increase in %A_fast_ further out of the experimental range. Moreover, the rates of the slow desensitization components and the amplitudes of the fast components are both underestimated for C^4^ and C^5^, as well as for multiple mutant combinations (Supplementary Figure 5B & C).

### Model III: adding inter-subunit coupling provides the best fit to experimental data

We finally improved the model by adding a degree of structural coupling between adjacent subunits. We postulated that desensitization of a particular subunit would favour desensitization of its neighbouring subunits. We thus incorporated coupling constants between subunit pairs in model II-β, and performed iterative kinetic modelling to minimize the divergences with electrophysiological data. The best fit was achieved assuming that, first, desensitization of SU4 decreases the recovery rate of SU5 by ε=10-fold - and vice versa, and second that desensitization of SU4 increases the desensitization rate of SU3 by γ=100-fold – and vice versa (Figure 6A, Supplementary Table 3). Apart from these couplings, model III retains all features from model II-β (Figure 6A&B). Of note, we don’t need to include any effect of SU4 or SU5 mutation on the recovery from desensitization (i.e. c_4_^-^ = c_5_^-^ = 1; Supplementary Table 3).

**Figure 6.**
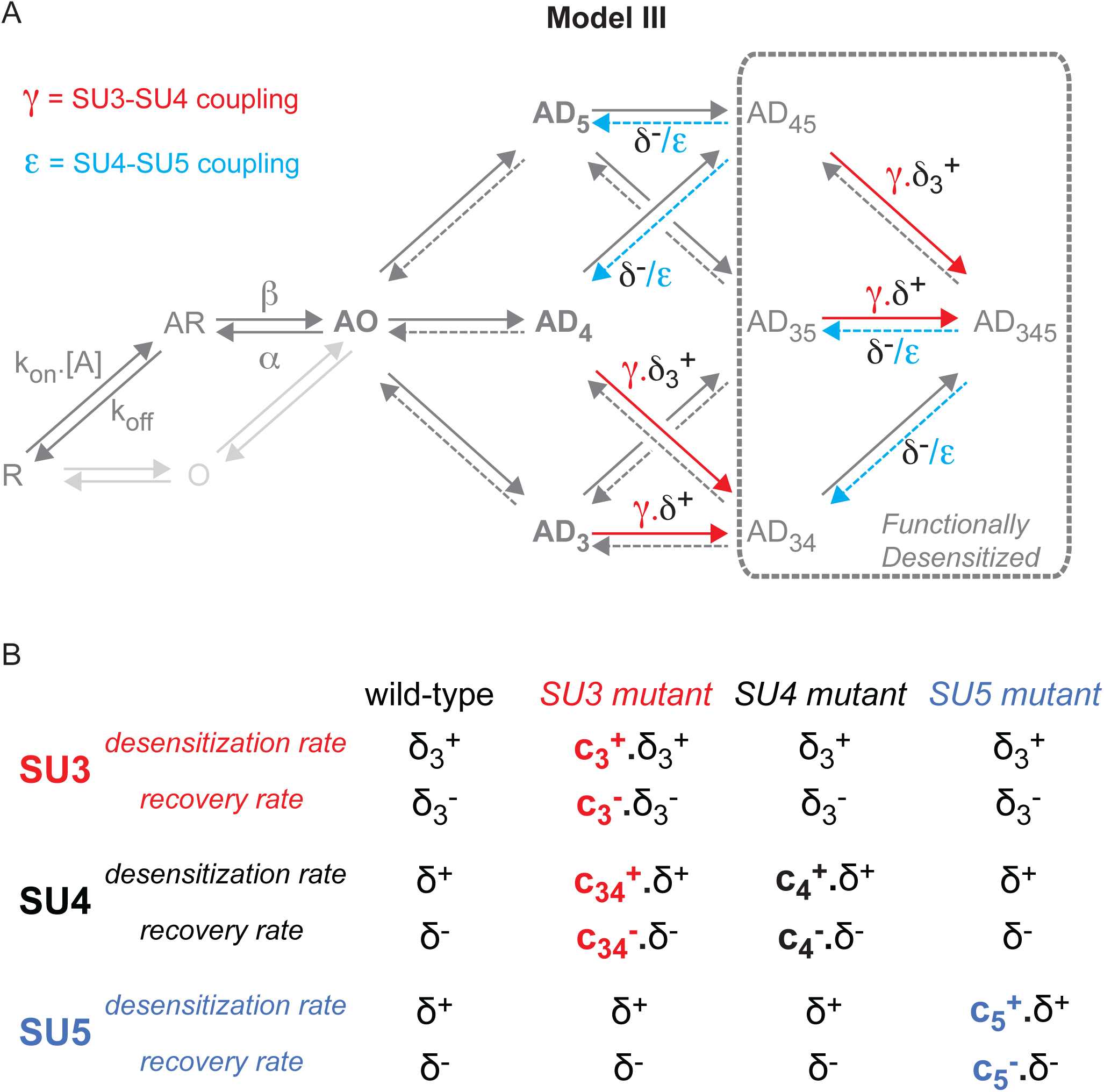
Model III introduces inter-subunit coupling during desensitization. ***A***, For the wild-type receptors, Model III builds upon Model II by adding some coupling between adjacent subunits during desensitization. On the one hand, desensitization of SU3 accelerates the desensitization of SU4 by a factor γ, and reciprocally. On the other hand, desensitization of SU4 slows the recovery of SU5 by a factor ε, and reciprocally. ***B***, For mutated concatemers, Model III incorporates the additional hypothesis that the mutation of SU3 also affects the desensitization of SU4 by increasing both its desensitization and recovery rates, by ratios c_34_^+^ and c_34_^-^, respectively.

As shown in Figure 7, model III largely accounts for experimental data, with experimental traces and simulated responses overlaying well (Figure 7A-H), including for the wild-type situation. The kinetics of the fast desensitization component, which are the most reliable experimental constraints in the dataset, are particularly well simulated (Figure 7I). The amplitudes of the fast component are overall in good agreement with the data, even though they are significantly underestimated for constructs C^5^, C^34^ and C^35^ (Figure 7K), while slow desensitization rates and steady-state currents are also underestimated for C^45^ and C^345^ (Figure 7J & L). Those minor discrepancies might reflect the contribution of SU1 and/or SU2 to the receptors’ desensitization (see Discussion below). Measurement errors on residual currents and slow desensitization kinetics might also occur when recording strongly desensitizing constructs.

**Figure 7.**
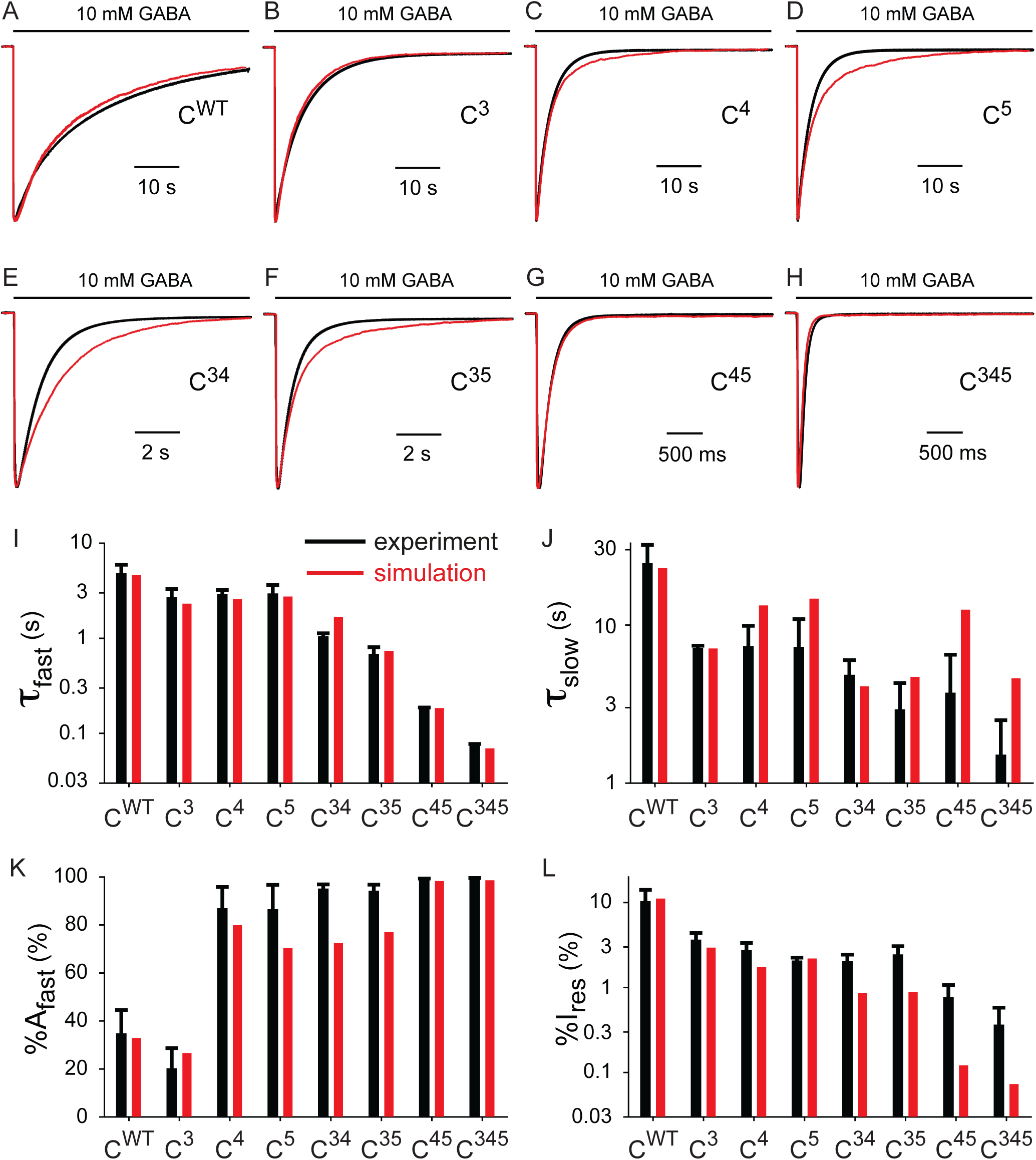
Model III simulations are broadly consistent with experimental data. ***A-H***, Representative currents for the indicated constructs, in black, are overlaid with their simulation counterparts in red. Note the changes in timescales. ***I-L***, Bar graphs summarizing the experimental data (in black) vs the simulations (in red) for the indicated concatemers on the kinetics of the fast (panel ***I***) and slow (panel ***J***) desensitization components, the relative amplitude of the fast component (panel ***K***) and the residual current after a 1 min long application of 10 mM GABA (panel ***L***). See Supplementary Table 3 for the numerical values of parameters.

Altogether, the whole set of data is consistent with a non-concerted model for pLGICs’ desensitization, characterized by three main features: 1) subunits can rearrange one at a time during desensitization, the multiple temporal components of desensitization reflecting the existence of intermediate asymmetrical desensitized states; 2) rearrangements of adjacent subunits during desensitization are nonetheless partially coupled; and 3) the desensitization of at least two subunits is required to shut the pore, i.e. to lead to functional desensitization.

## DISCUSSION

To illustrate the main features of our model of wild-type α1β2γ2 GABA_A_Rs desensitization, we show in Figure 8A the time-dependence of the various desensitized states’ occupancies during desensitization. Since SU4 and SU5 desensitize the fastest, the receptors in the active state will transit first through a pre-desensitized open-pore state, in which either SU4 or SU5 is desensitized (Supplementary Figure 6). Functional desensitization, i.e. loss of electrophysiological response, subsequently occurs upon desensitization of the second fast subunit to yield the AD_45_ state (Figure 8A). The final step along the desensitization pathway would correspond to the desensitization of SU3, resulting in the slow component of desensitization, i.e. the entry in the AD_345_ state (Figure 8A). Like in all kinetic schemes where the slow- and fast-desensitized states are connected, this final step slowly depletes receptors from the fast-desensitized pool, which in turn displaces the overall population away from active conformations. We can thus extract the kinetically favoured pathway and provide a schematic structural depiction of the movements of the M2 helices during desensitization, as shown in Figure 8B. Interestingly, the requirement for two desensitized subunits to occlude the pore provides a framework to interpret results at α7 nAChRs, whose desensitization is blocked by PNU-120596. Indeed, it was shown that at least four α7 subunits need to be bound by PNU-120596 in order to block desensitization, meaning that as soon as two subunits are unbound, the receptors can undergo functional desensitization^22^.

**Figure 8.**
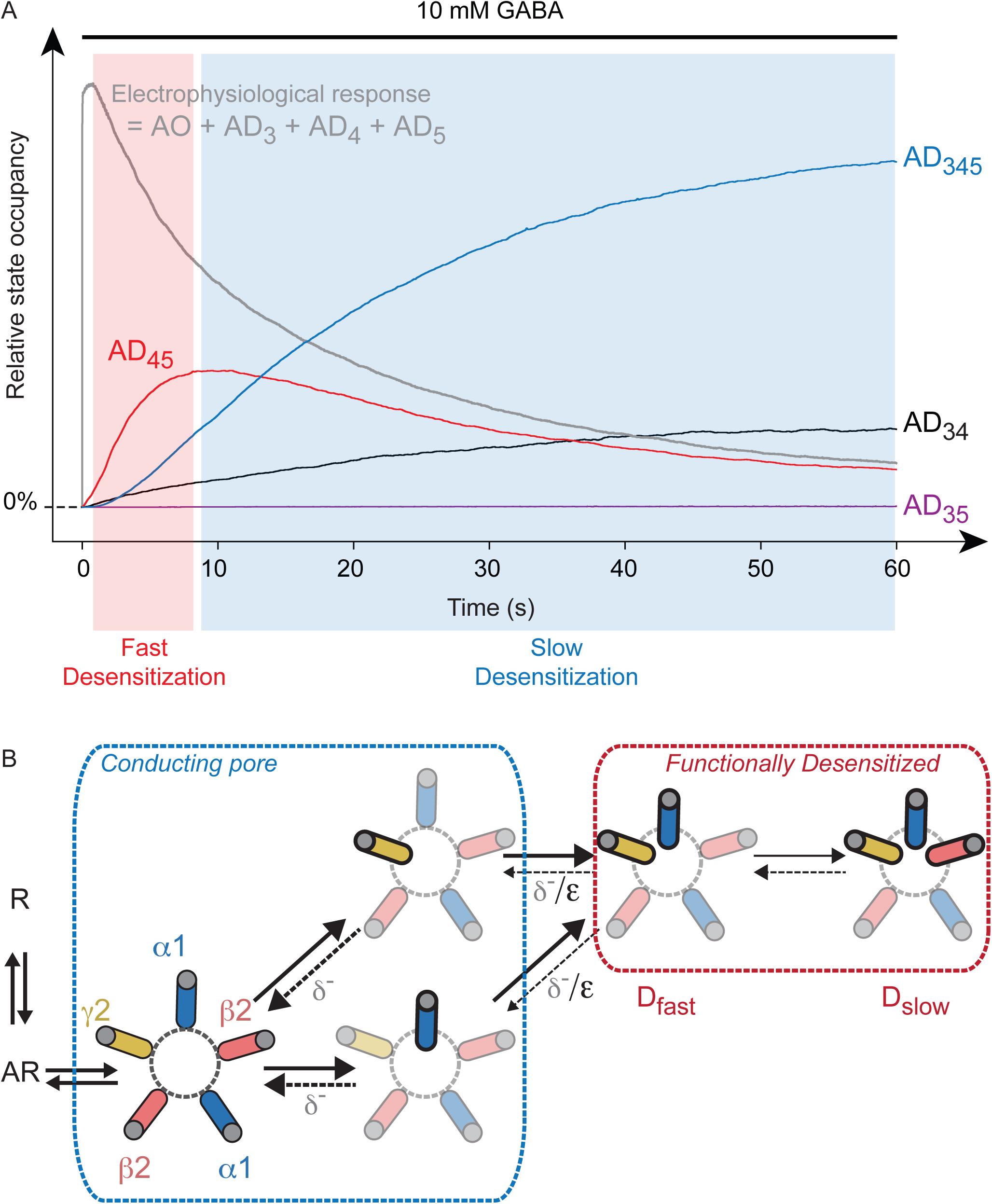
States occupancies predictions and structural depiction of Model III. ***A***, The overall population of wild-type receptors in an active conformation is compared to the relative occupancies of the various desensitized states. As depicted by the red box, the early phase of desensitization is carried by the AD_45_ state. On longer time scales (blue box), slow desensitization is largely embodied by the entry in the AD_345_ state. The analysis of states occupancies was performed with QuB simulations. ***B***, In this simplified depiction of model III, we extracted the kinetically favoured pathway for the desensitization of wild-type α1β2γ2 GABA_A_Rs. Upon agonist binding, the receptor is transiently stabilized in a fully open pseudo-symmetrical conformation. The two first subunits to rearrange during desensitization are the α1 and the γ2 subunits involved in the binding of benzodiazepines, namely SU4 and SU5 in our concatemers. While one desensitized subunit is not enough to occlude the pore, fast desensitization corresponds to the rearrangement of both SU4 and SU5 subunits, which are coupled. Slow desensitization is then driven by the slower rearrangement of the SU3 subunit, i.e. the β2 subunit opposite to the γ2 subunit.

The whole set of data points to the γ2 subunit as a major determinant of the desensitization of α1β2γ2 GABA_A_Rs. Interestingly, the TMD of the γ2 subunit appears highly flexible in detergent conditions, its TMD collapsing within the pore when α1β3γ2 GABA_A_Rs are solubilized in decylmaltoside neopentylglycol^23^ or n-dodecyl-β-D-maltopyranoside^24,25^. The addition of lipids stabilizes the γ2 TMD in a more physiologically relevant conformation^23^, but it still remains highly mobile and necessitates nanodiscs to be well resolved^17^. While the lack of the M3-M4 intracellular loop might impact the structures solved in detergent, it is tempting to speculate that the dynamic nature of the γ2 TMD during desensitization is a functional counterpart of this biochemical instability. It is also interesting to note that the γ2 subunit contains a phosphorylation site at a serine located at the intracellular end of the M3 segment, namely S327^26^. This residue is located in an intracellular cassette modulating the desensitization properties of inhibitory pLGICs^8^, only eight residues downstream of the M3-5’ residues that we have targeted in the current study. One could thus easily imagine that phosphorylation of γ2-S327 provides a mean to modulate the desensitization of γ2-containing GABA_A_Rs. Last but not least, the prominent role of the γ2 subunit in shaping the desensitization of α1β2γ2 GABA_A_Rs makes it an ideal target for pharmacological modulation of these receptors. Modulating desensitization should barely affect basic synaptic signalling, which could thus lead to fairly safe compounds with a large therapeutic window. Targeting the γ2 subunit specifically, in a desensitization locus with divergent sequences among pLGICs such as the intracellular end of the M3 segment, should also provide an easy mean to achieve subtype selectivity.

The apparent lack of effect on desensitization when mutating SU1 or SU2 alone is another striking feature of the dataset. A first hypothesis might be that these subunits don’t desensitize during the one-minute-long GABA application from our protocol. This is unlikely: in that case, mutating SU1 and/or SU2 should not affect the fast desensitization of concatemers harbouring mutations on other subunits. However, mutating SU1 and SU2 leads to an almost 2-fold increase in the fast desensitization kinetics of C^345^ (Figure 2; Supplementary Table 1). An alternative hypothesis would then be that SU1 and/or SU2 display very fast desensitization, but with a desensitization equilibrium largely displaced towards their open conformation (“δ^+^/ δ^-^ << 1”), thereby barely contributing to the macroscopic course of desensitization. Such desensitization equilibrium would minimally affect the size of currents, nor the apparent affinity for the agonist. In that event, it is conceivable that SU1 and/or SU2 mutations effects could be revealed on a mutant background owing to inter-subunit coupling. This potential impact of inter-subunit coupling involving SU1 or SU2 might also explain why our kinetic simulations slightly differ from experiments for certain mutants. Our dataset unfortunately provides too little constraint to build a comprehensive scheme for the contribution of SU1 and SU2, and prevents their inclusion in our kinetic model.

Our non-concerted asymmetrical model provides a clear departure from a classical view in which D_fast_ and D_slow_ states are fundamentally different. It raises the possibility that these states are identical at the single-subunit level, with D_fast_ only reflecting asymmetrical intermediates, mainly AD_45_, along the desensitization process. Such scheme might appear surprising given the widely accepted concerted nature of pLGICs gating, as described for the muscle-type nAChR^27^. However, the analysis and concepts in favour of a concerted gating of pLGICs, like the MWC model framed more than half a century ago^28^, have largely focused on biochemical and electrophysiological data obtained under gating equilibrium conditions such as concentration-response curves^21,27^. In the case of desensitization, the events are slow enough that intermediate events are directly detectable, namely the D_fast_ state(s). If one could record the activation kinetics with sufficient temporal precision, it is likely that proper data fitting would also require the use of non-concerted asymmetric rearrangements. This is actually hinted by the prime model of muscle-type nAChR activation, in which conformational changes can affect independently either of the two ACh binding sites^29^, as well as by rate-equilibrium free energy relationship analyses arguing for non-concerted rearrangements of M2 helices during nAChR activation^30^. Moreover, molecular dynamics studies also pinpoint the cytoplasmic end of the pore as a locus for asymmetric conformations at the µs timescale: the five −2’ residues are often distributed in a non-symmetrical fashion during simulations of the open state of the zebrafish α1 Glycine receptor^31,32^. Of note, channels and receptors from other families are also known to rely on asymmetric gating. This is the case of the prokaryotic magnesium channel CorA, whose active state actually stems from an asymmetric conformation as reported by cryo-electron microscopy^33^. This is also the case for NMDA receptors, for which the cryo-electron microscopy of tri-heteromeric GluN1/GluN2A/GluN2B receptors reveals an asymmetric organisation^34^.

The exact structural underpinnings of desensitization remain however ill-defined, in particular since the current structures have been obtained for presumable resting and desensitized conformations so far^10,17^. In the absence of an active conformation, one can only speculate on the precise molecular events occurring during the active to desensitized transition.

## METHODS

### Molecular biology

The GABA_A_ concatemeric α1β2γ2 construct was previously described^18^., based on the concatenation of mouse GABA_A_ subunits. Briefly, the five subunits were subcloned in the order β2-α1-β2-α1-γ2 into a low copy number vector pRK5, retaining the peptide signal of the first subunit only. We used the short splice variant of the γ2 subunit, γ2S. All five subunits are flanked by unique restriction sites to allow the subcloning of mutated subunits, and separated by 15-20 residues long polyglutamine linkers, depending on the length of the C-terminus end of the subunit preceding the linker. The construct thus shows the arrangement ClaI-β2-20Q-AgeI-α1-15G-SalI-β2-20Q-NheI-α1-15Q-γ2S-Stop-HindIII. Site-directed mutagenesis was performed on individual subunits as previously described^8^. Owing to the unique restriction sites, mutated subunits where then sequentially subcloned in the concatemer to yield the desired combinations of mutated subunits. We finally sequenced the resulting mutated concatemers to check for the incorporation of the desired mutated subunits. We could not use primers annealing anywhere in α1 or β2 for sequencing, as both subunits are present as duplicates in the concatemer. Instead, we sequenced SU1-4 subunits with primers annealing at their 5’ DNA extremity, centered on the sequence of the unique restriction site preceding the following subunit. Such reverse primers enable the sequencing of the 5’ end of the subunits’ DNA, coding for their C-terminus once translated.

### Expressing GABA_A_Rs in Xenopus laevis oocytes

Ovaries from *Xenopus laevis* were obtained from CRB Xenopes in Rennes. Free oocytes were obtained by incubating segments of ovary in collagenase type 1 (Sigma) dissolved in a Ca^2+^-free OR2 solution, which contained (mM): 85 NaCl, 5 HEPES, 1 MgCl_2_, pH adjusted to 7.6 with KOH. After 2-4 hrs exposure to collagenase I, defolliculated oocytes were washed several times with OR2, and thereafter maintained in a Barth’s solution containing (mM): 88 NaCl, 1 KCl, 0.33 Ca(NO_3_)_2_, 0.41 CaCl_2_, 0.82 MgSO_4_, 2.4 NaHCO_3_, 10 HEPES, pH adjusted to 7.6 with NaOH. Single oocytes were injected with 27.6 nl of concatemeric GABA_A_R cDNAs (nuclear injection) at a concentration of 30 ng/µl. Oocytes were incubated at 17°C in Barth’s solution devoid of serum or antibiotics.

### Two-electrode voltage clamp recording

Oocytes expressing pentameric concatemers were recorded 2-4 days after injection. They were superfused with a solution containing (mM): 100 NaCl, 2 KCl, 2 CaCl_2_, 1 MgCl_2_, 5 HEPES, pH adjusted to 7.4 with NaOH. Solution flowed at an approximate speed of 12 mL/min. Currents were recorded using a Warner OC-725C amplifier, a Digidata 1550A interface and pCLAMP 10 (Molecular Devices). Currents were digitized at 500 Hz and filtered at 100 Hz (30-60 Hz used for display purposes). Oocytes were voltage-clamped at −60 mV and experiments conducted at room temperature. Desensitising currents were induced by 1 min applications of 10 mM GABA.

### Data analysis

The extent of desensitisation was determined as (1 – I_res_/I_peak_), where I_peak_ is the peak current and I_res_ the residual current remaining at the end of the agonist application. Weighted decay time constants for desensitisation were determined by fitting the desensitising phase with two exponential components (pCLAMP 10), as given by the following equation: τ_w_ = %A_fast_ * τ_fast_ + (1-%A_fast_) * τ_slow_. All data values are means ± standard deviation.

### Drugs and chemicals

All compounds were purchased from Sigma. GABA was prepared as a 1 M stock solution in recording solution. Aliquots were stored at −20°C.

### Kinetic modelling

We used QUB^20^ to build Markov-chain kinetic models. Each simulation contained 10,000 – 30,000 channels. The binding and gating rate constants are broadly consistent with previously published values for GABA_A_Rs^35^. For each model, we performed iterative rounds of kinetic simulations by adjusting manually the set of parameters. Binding and gating constants being fixed, Model I (Figure 4), Model II (Figure 5), Model II-β (Supplementary Figure 4) and the concerted model (Supplementary Figure 2) only contain four parameters for the wild-type receptors (δ^+^, δ^-^, δ_3_^+^ and δ_3_^-^ for Models I, II and II-β; fast and slow desensitization rates and their recovery counterparts for the concerted model). This equates to the number of independent experimental measurements related to the two desensitization components (τ_fast_, τ_slow_, %A_fast_, %I_res_). We could thus be confident that, once we have a set of parameters accounting for the wild-type data, the model has a good predictive value. Mutation-induced changes in those parameters (c_3_^+^, c_3_^-^, c_4_^+^, c_4_^-^, c_5_^+^, c_5_^-^) for models Models I, II were then adjusted manually to account for the effects of individual mutants (C^3^, C^4^, and C^5^), Model II-β requiring the additional adjustment of c_34_^+^, c_34_^-^ for the effect of SU3 mutation. The effects of mutations in the concerted model (Supplementary Figure 2) led to four mutation-related parameters (γ_f_, ε_f_, γ_s_ and ε_s_) for each individual mutant C^3^, C^4^, and C^5^. In all cases, mutation-related parameters derived from individual mutants were then combined to predict the effect of combining and mutations in constructs C^34^, C^35^, C^45^ and C^345^. For Model III, we generated a series of wild-type models with values for coupling constants (γ and ε) in the 1-1000 range (1, 10, 100 and 1000). We next manually adjusted the mutation-induced changes as described above for Model II-β.

## Supporting information

Supplementary discussion, data and tables

## Abbreviations

ECD: extracellular domain;
GABA: γ-aminobutyric acid;
GABA_A_R: type A γ-aminobutyric acid receptor;
pLGIC: pentameric ligand-gated ion channel;
MWC: Monod-Wyman-Changeux;
nAChR: nicotinic acetylcholine receptor;
TEVC: two-electrode voltage clamp;
TMD: transmembrane domain.

## Acknowledgements

The authors would like to thank Drs Thomas Boulin, Hugues Nury, Laurie Peverini, Marie Prevost and Prof Trevor Smart for critical reading of the manuscript; and acknowledge financial support by the Fondation de la Recherche Médicale (grant “Équipe FRM” DEQ20140329497 to P-J.C.) and the European Commission Research Executive Agency (Marie Sklodowska-Curie Action, Individual Fellowship 659371 to M.G.).

