## Supplementary discussion, data and tables for "The desensitization pathway of GABA_A_ receptors, one subunit at a time"

#### Model III: limitations and implications for single-channel recordings

We have to acknowledge here two main shortfalls of our model. First, no unliganded desensitized state is present, although these are known to exist and contribute to the recovery from desensitization upon removal of the agonist[1]. Our kinetic model thus cannot be used to model recovery from desensitization. Second, at the single channel level, our model makes a strong prediction when the channel transits from the open state to a more stable pre-desensitized state still capable of conducting ions. Indeed, in the open state, single-channel openings on the ms timescale are separated by brief shut times reflecting sojourns in resting or pre-active states[2], giving rise to a maximum open probability of about 70%-80% for α1β2γ2 $GABA_A$Rs[3]. However, once the channel enters a pre-desensitized state in our final scheme – i.e., $AD_3$, $AD_4$ or $AD_5$ -, the predicted lifetime of such state should produce uninterrupted single-channel openings of several seconds for wild-type receptors. Such openings are not observed in single-channel recordings. It is likely that, when one subunit enters its desensitized conformation, the other subunits retain the ability to visit their resting conformation, thereby providing a way to maintain an open probability well below unity. Still, one could expect a strong influence of the receptor's desensitization on its single-channel gating efficacy under equilibrium conditions, e.g. in cell-attached patch-clamp recordings. It is thus striking that a recent study highlighted such phenomenon at α1 glycine receptors (GlyRs)[4]. Indeed, Ivica *et al.* showed that deletion of the intracellular domain (ICD) of homomeric human α1 GlyRs largely increases their efficacy of gating. At the same time, the ICD is known to affect the desensitization of α1 GlyRs[5–7], and the study by Ivica *et al.* displays single-channel clusters that seem largely lengthened by the ICD deletion, suggesting that their ICD-deleted construct shows reduced levels of desensitization (comparing the glycine and alanine recordings from panels from Figure 2B and Figure 3B in ref[4]). It is thus tempting to speculate that the ICD-deleted GlyRs constructs from the study by Ivica *et al.* show a greater stability of their pre-desensitized states with one single subunit being desensitized, resulting in both lengthened single-channel clusters and higher cluster open probability under equilibrium conditions.

Our model makes another prediction regarding single-channel recordings, which would provide strikingly different estimation of desensitization kinetics under equilibrium versus non-equilibrium conditions. Indeed, with cell-attached recordings performed in the continued presence of the agonist, some desensitized channels would spontaneously recover and visit a conducting state, amenable to electrophysiological detection. Given our set of parameters, the kinetically favoured pathway would be for a receptor in a $AD_{345}$ state to transit to the $AD_{34}$ state and then to the conducting pre-desensitized $AD_3$ state. From there, the favoured transition would be to re-enter the $AD_{34}$ state with a microscopic rate of $\gamma.\delta^+$, i.e. 28 $s^{-1}$, resulting in a cluster duration of about 36 ms. On the contrary, a receptor recorded in non-equilibrium conditions, such as performed in fast-perfusion experiments in the outside-out configuration or in whole-cell experiments, would start off in the resting state R, transit to the agonist-bound AR state, and then finally open (AO state). The electrophysiological current would be detectable for the entire duration from the entry to the open state to the exit in a fully desensitized state. In this case, the favoured routes would either be AO to $AD_4$ to $AD_{45}$ or AO to $AD_5$ to $AD_{45}$. The apparent corresponding macroscopic rate would roughly equate to $2.(\delta^+/\delta^-).\delta^+$, i.e. about 0.25 $s^{-1}$, resulting in a cluster duration of about 4 s. Thus, our model suggests that cell-attached patch-clamp recordings might not be predictive of the desensitization properties of $\alpha1\beta2\gamma2$ $GABA_ARs$ driven by a sudden rise in agonist concentration, as observed during phasic synaptic release.

### SUPPLEMENTARY DATA

**Supplementary Figure 1. Relative effect of M3-5' mutations on the weighted desensitization kinetics of a series of background mutated concatemers.** Each bar plot corresponds to the relative effect of the M3-5' mutation of the indicated subunit on the weighted desensitization rate of the background constructs.

**Supplementary Figure 2. A concerted model only including pseudo-symmetrical states cannot account for the synergistic effects of SU4 and SU5 M3-5' valine mutations.** ***A-D***, Simplified concerted kinetic schemes used to model wild-type (panel ***a***), single mutant (panels ***B*** & ***C***) and double mutant (panel ***D***) concatemers. While unliganded openings are possible at all ligand-gated receptors, consistent with a Monod-Wyman-Changeux model of allosteric proteins, they are exceedingly rare at wild-type α1β2γ2 $GABA_A$Rs (see Main Text), and we therefore decided not to include them in our kinetic modelling. In the absence of direct interaction between the mutated residues, mutations are expected to have additive effects (see Main Text) as described in panel ***D***. ***E***, Putative free energy diagram translating the kinetic schemes from panels ***A-D***. ***F-G***, Representative currents (in black) from $C^{WT}$, $C^4$, $C^5$ and $C^{45}$ constructs are overlaid with the simulated $C^{45}$ response (in red) as predicted from the concerted model adjusted to account for the wild-type and single mutant constructs ($C^{WT}$, $C^4$ and $C^5$). Note that the concerted model largely underestimates the desensitization kinetics of the double mutant $C^{45}$.

**Supplementary Figure 3. Simulations from Model II.** ***A-D***, Bar graphs summarizing the experimental data (in black) vs the simulations (in red) for the indicated concatemers on the kinetics of the fast (panel ***A***) and slow (panel ***B***) desensitization components, the relative amplitude of the fast component (panel ***C***) and the residual current after a 1 min long application of 10 mM GABA (panel ***D***). See Supplementary Table 3 for the numerical values of parameters.

**Supplementary Figure 4. Model II-β: mutation of SU3 directly impacts on SU4 desensitization.** ***A***, For wild-type receptors, Model II and Model II-β are strictly identical. ***B***, For mutated concatemers, Model II-β incorporates the hypothesis that the mutation of SU3 also affects the desensitization of SU4 by increasing both its desensitization and recovery rates, by ratios $c_{34}^{+}$ and $c_{34}^{-}$, respectively.

**Supplementary Figure 5. Simulations from Model II-β.** ***A-D***, Bar graphs summarizing the experimental data (in black) vs the simulations (in red) for the indicated concatemers on the kinetics of the fast (panel ***A***) and slow (panel ***B***) desensitization components, the relative amplitude of the fast component (panel ***C***) and the residual current after a 1 min long application of 10 mM GABA (panel ***D***). See Supplementary Table 3 for the numerical values of parameters. ***E***, Overlay of a representative current for $C^{WT}$ (in black) with the response simulated according to Model II-β (in red). Note that both the amplitude of the fast desensitization component and residual currents are overestimated in the simulation.

**Supplementary Figure 6. Time-dependant changes in the occupancy of the receptors's states as predicted by model III.** The relative occupancies of each state contributing to the electrophysiological currents, namely AO, $AD_3$, $AD_4$ and $AD_5$, are compared to the overall population of receptors in an active conformation ($AO + AD_3 + AD_4 + AD_5$), which directly translates the experimental recordings. The analysis of states occupancies was performed with QuB simulations.

**Supplementary Table 1. Kinetics and extent of desensitization of the pentameric $GABA_A$ concatemers used in the present study.** All data are shown as means ± standard deviations. For each construct, the number of *Xenopus laevis* oocytes that we recorded from is indicated on the right. At least two different batches of oocytes were used for all constructs.

**Supplementary Table 2. Numerical values used for the parameters from the concerted model in Supplementary Figure 1.**

**Supplementary Table 3. Numerical values used for the parameters from the non concerted models I, II, II-β and III.**

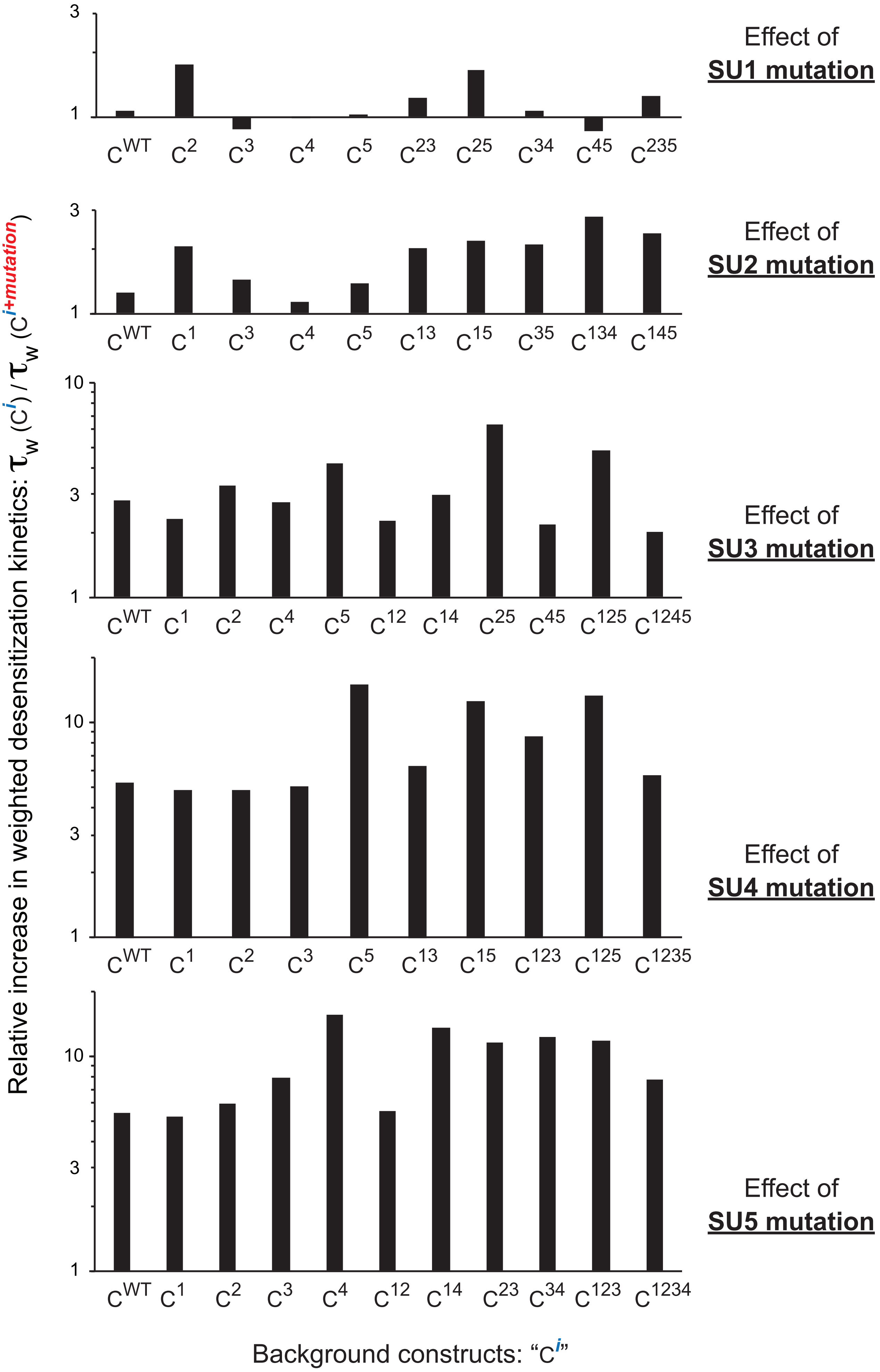


**Supplementary Figure 1**

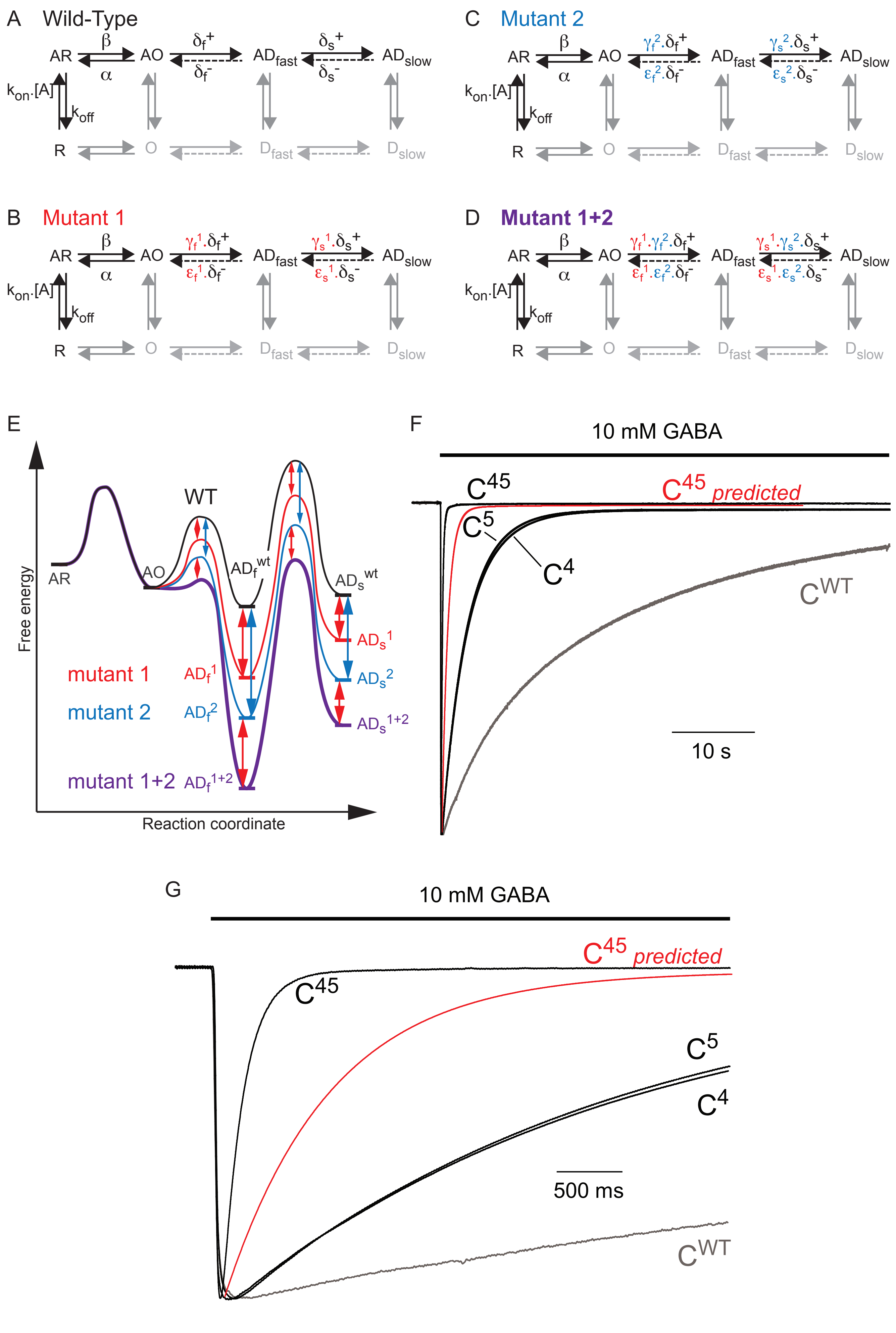


**Supplementary Figure 2**

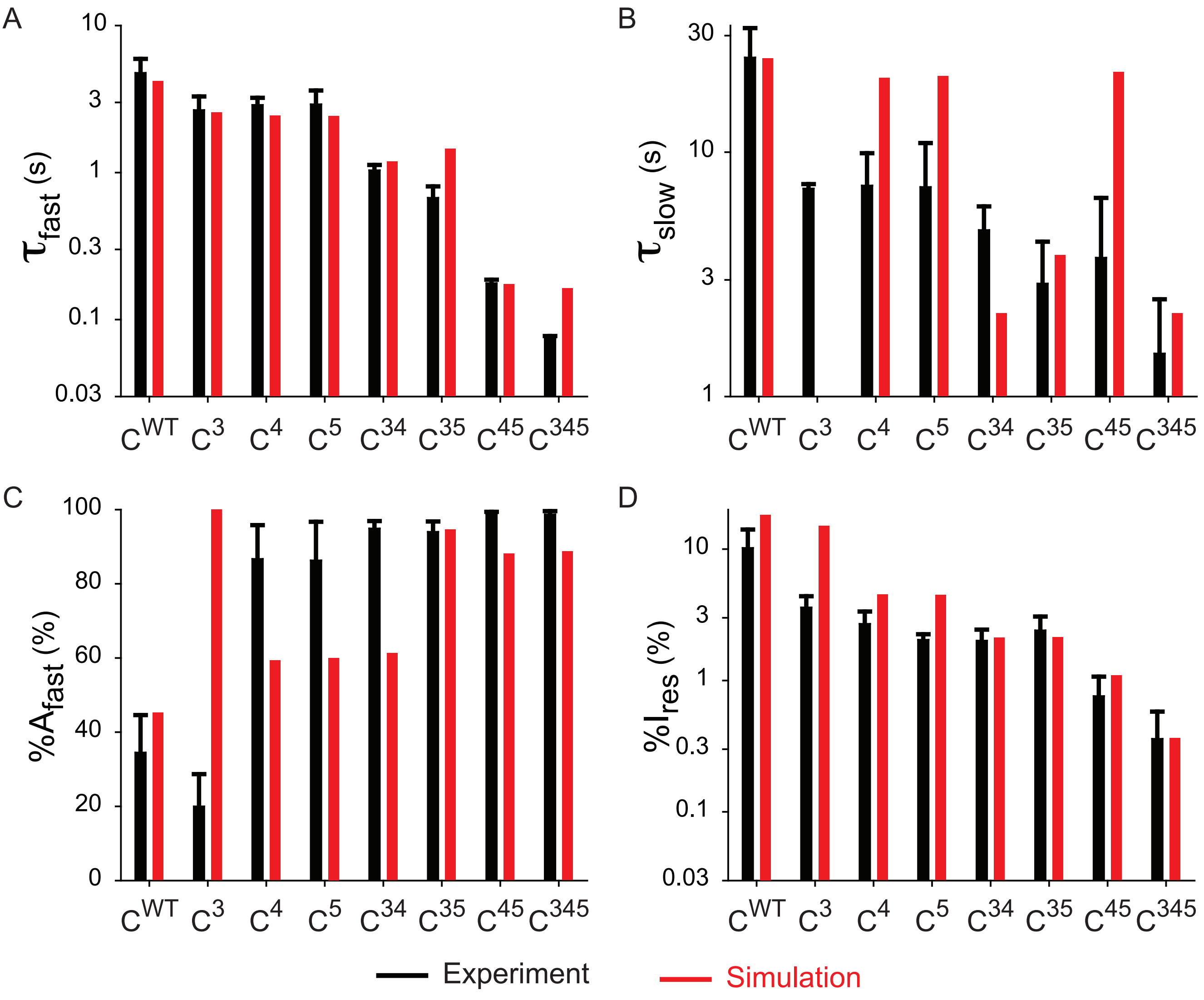


**Model II**

**Supplementary Figure 3**

A

**Model II-β**

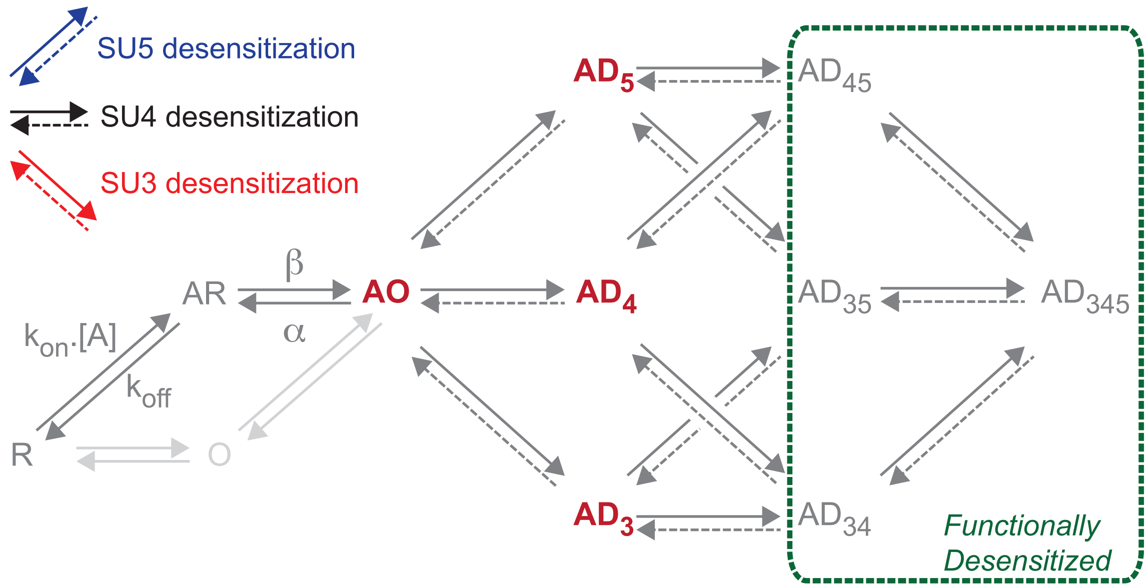


B

| | | wild-type | SU3 mutant | SU4 mutant | SU5 mutant |
|---|---|---|---|---|---|
| **SU3** | *desensitization rate* | $\delta_3^+$ | $c_3^+.\delta_3^+$ | $\delta_3^+$ | $\delta_3^+$ |
| | *recovery rate* | $\delta_3^-$ | $c_3^-.\delta_3^-$ | $\delta_3^-$ | $\delta_3^-$ |
| **SU4** | *desensitization rate* | $\delta^+$ | $c_{34}^+.\delta^+$ | $c_4^+.\delta^+$ | $\delta^+$ |
| | *recovery rate* | $\delta^-$ | $c_{34}^-.\delta^-$ | $c_4^-.\delta^-$ | $\delta^-$ |
| **SU5** | *desensitization rate* | $\delta^+$ | $\delta^+$ | $\delta^+$ | $c_5^+.\delta^+$ |
| | *recovery rate* | $\delta^-$ | $\delta^-$ | $\delta^-$ | $c_5^-.\delta^-$ |

**Supplementary Figure 4**

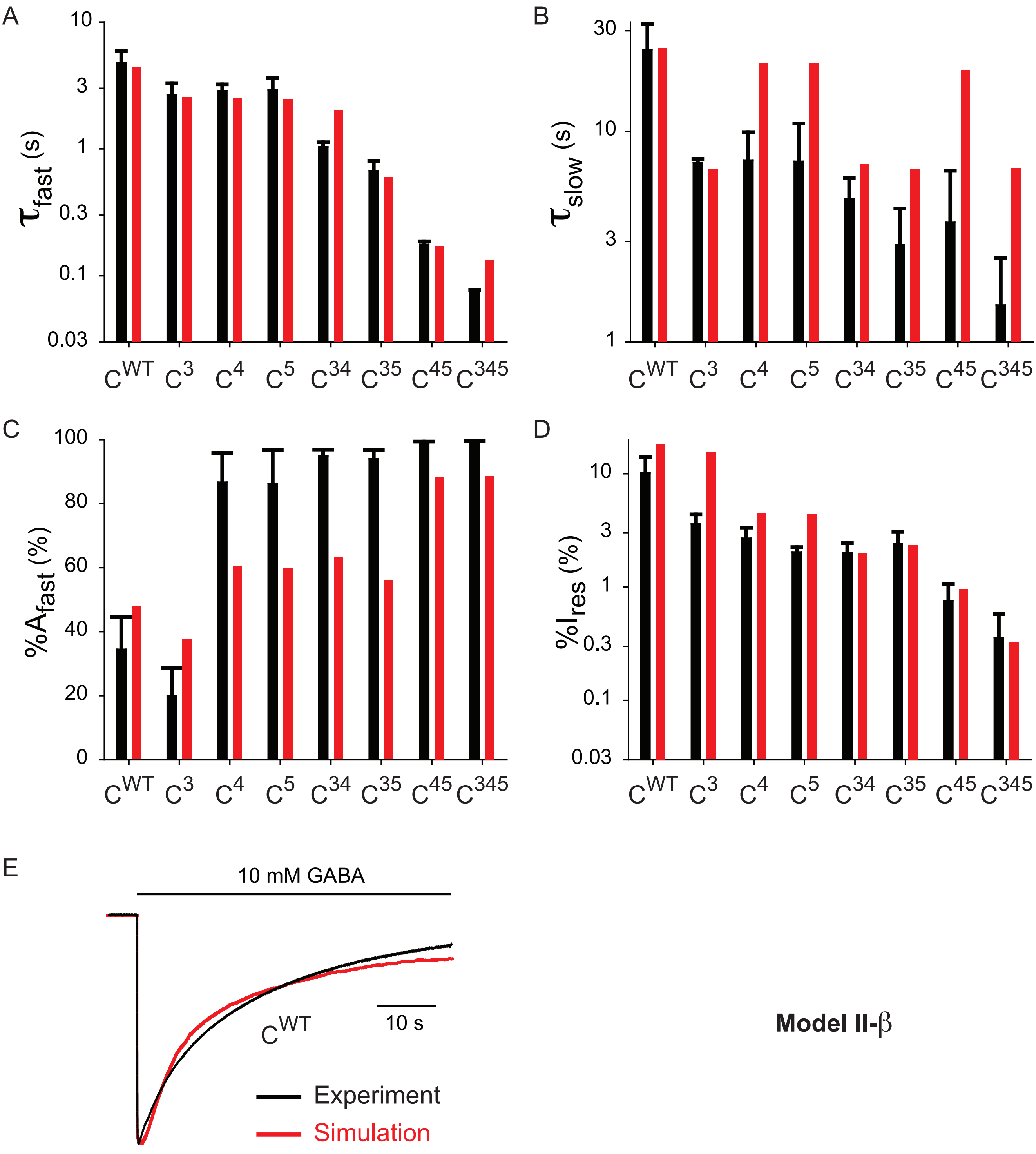


**Supplementary Figure 5**

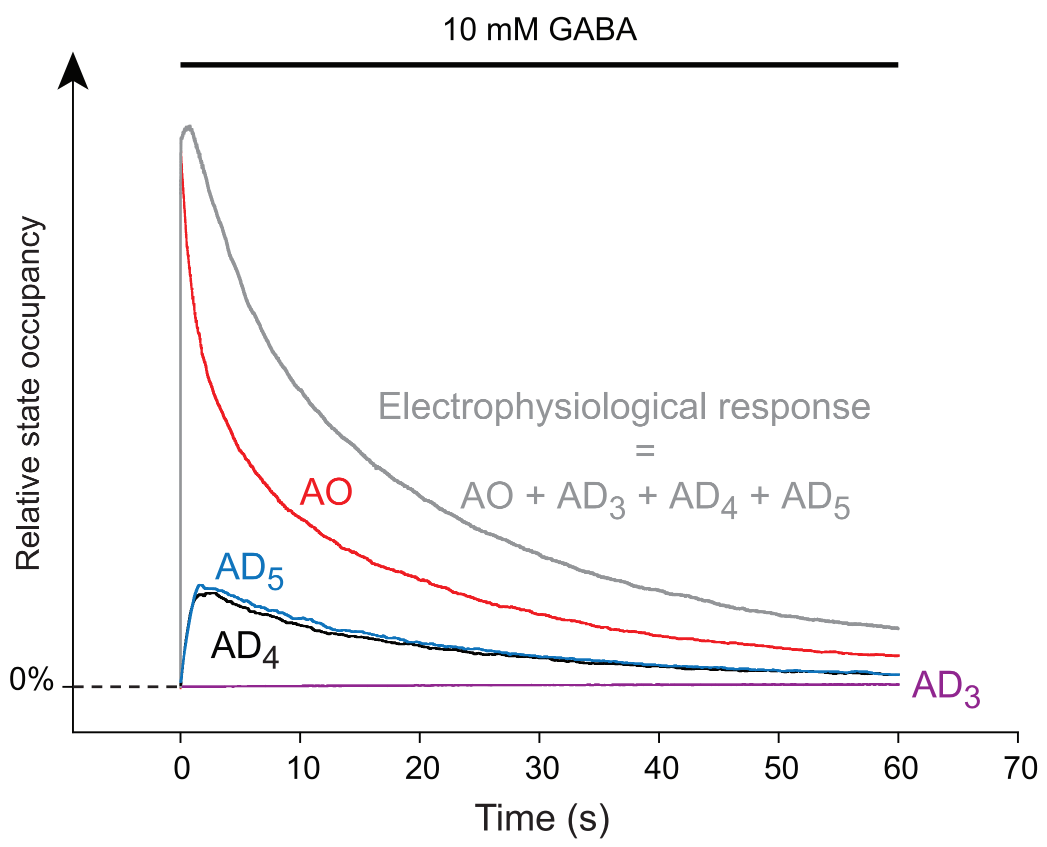


**Supplementary Figure 6**

| Construct | SU1 | SU2 | SU3 | SU4 | SU5 | $\tau_{fast}$ (s) | $\tau_{slow}$ (s) | %$A_{fast}$ | $\tau_w$ (s) | %$I_{res}$ | # cells |
|---|---|---|---|---|---|---|---|---|---|---|---|
| $C^{WT}$ | WT | WT | WT | WT | WT | 4,8 ± 1,2 | 24,4 ± 7,8 | 34,5% ± 10,1% | 17,6 ± 5,1 | 10,2% ± 3,9% | n = 40 |
| $C^{1}$ | mut | WT | WT | WT | WT | 4,5 ± 0,7 | 23,0 ± 7,0 | 35,9% ± 4,9% | 16,6 ± 5,7 | 9,1% ± 3,4% | n = 5 |
| $C^{2}$ | WT | mut | WT | WT | WT | 4,0 ± 0,9 | 18,5 ± 5,1 | 29,9% ± 5,5% | 14,2 ± 4,5 | 7,5% ± 2,9% | n = 5 |
| $C^{3}$ | WT | WT | mut | WT | WT | 2,7 ± 0,6 | 7,1 ± 0,3 | 20,0% ± 8,7% | 6,2 ± 0,4 | 3,6% ± 0,8% | n = 4 |
| $C^{4}$ | WT | WT | WT | mut | WT | 2,9 ± 0,3 | 7,3 ± 2,6 | 86,7% ± 9,1% | 3,4 ± 0,2 | 2,7% ± 0,7% | n = 5 |
| $C^{5}$ | WT | WT | WT | WT | mut | 2,9 ± 0,7 | 7,2 ± 3,7 | 86,3% ± 10,4% | 3,3 ± 0,8 | 2,0% ± 0,2% | n = 8 |
| $C^{12}$ | mut | mut | WT | WT | WT | 3,6 ± 0,6 | 10,7 ± 1,2 | 35,8% ± 8,1% | 8,1 ± 0,9 | 2,8% ± 0,2% | n = 10 |
| $C^{13}$ | mut | WT | mut | WT | WT | 3,7 ± 0,4 | 9,2 ± 0,9 | 36,9% ± 19,7% | 7,2 ± 1,6 | 3,5% ± 0,5% | n = 5 |
| $C^{14}$ | mut | WT | WT | mut | WT | 3,3 ± 0,3 | 4,2 ± 1,2 | 60,8% ± 18,6% | 3,4 ± 0,2 | 2,7% ± 1,0% | n = 7 |
| $C^{15}$ | mut | WT | WT | WT | mut | 2,7 ± 0,9 | 9,7 ± 4,4 | 83,4% ± 32,1% | 3,2 ± 0,9 | 2,5% ± 0,6% | n = 6 |
| $C^{23}$ | WT | mut | mut | WT | WT | 2,8 ± 0,3 | 5,7 ± 0,6 | 45,6% ± 18,2% | 4,3 ± 0,6 | 3,3% ± 0,3% | n = 6 |
| $C^{24}$ | WT | mut | WT | mut | WT | 2,7 ± 0,4 | 12,0 ± 5,8 | 95,7% ± 5,5% | 3,0 ± 0,4 | 2,2% ± 0,6% | n = 8 |
| $C^{25}$ | WT | mut | WT | WT | mut | 2,0 ± 0,7 | 5,7 ± 5,1 | 80,8% ± 25,0% | 2,4 ± 0,7 | 1,9% ± 0,8% | n = 7 |
| $C^{34}$ | WT | WT | mut | mut | WT | 1,0 ± 0,1 | 4,8 ± 1,2 | 94,9% ± 2,0% | 1,2 ± 0,1 | 2,0% ± 0,4% | n = 9 |
| $C^{35}$ | WT | WT | mut | WT | mut | 0,67 ± 0,13 | 2,9 ± 1,4 | 94,0% ± 2,8% | 0,79 ± 0,15 | 2,4% ± 0,7% | n = 14 |
| $C^{45}$ | WT | WT | WT | mut | mut | 0,18 ± 0,01 | 3,7 ± 2,8 | 98,8% ± 0,6% | 0,22 ± 0,03 | 0,8% ± 0,3% | n = 5 |
| $C^{123}$ | mut | mut | mut | WT | WT | 2,7 ± 0,8 | 6,8 ± 2,1 | 71,0% ± 20,1% | 3,6 ± 1,2 | 3,6% ± 1,1% | n = 7 |
| $C^{125}$ | mut | mut | WT | WT | mut | 1,3 ± 0,1 | 6,2 ± 1,6 | 96,1% ± 1,8% | 1,5 ± 0,1 | 1,7% ± 0,4% | n = 8 |
| $C^{134}$ | mut | WT | mut | mut | WT | 0,96 ± 0,13 | 3,9 ± 1,7 | 92,1% ± 3,5% | 1,2 ± 0,2 | 2,5% ± 0,7% | n = 7 |
| $C^{145}$ | mut | WT | WT | mut | mut | 0,16 ± 0,02 | 4,6 ± 2,7 | 98,0% ± 0,7% | 0,26 ± 0,09 | 0,6% ± 0,3% | n = 8 |
| $C^{235}$ | WT | mut | mut | WT | mut | 0,33 ± 0,05 | 3,2 ± 2,5 | 97,8% ± 1,2% | 0,38 ± 0,06 | 1,1% ± 0,4% | n = 6 |
| $C^{345}$ | WT | WT | mut | mut | mut | 0,07 ± 0,003 | 1,5 ± 1,0 | 98,7% ± 0,9% | 0,10 ± 0,04 | 0,4% ± 0,2% | n = 6 |
| $C^{1234}$ | mut | mut | mut | mut | WT | 0,36 ± 0,057 | 2,5 ± 1,9 | 96,0% ± 3,6% | 0,41 ± 0,07 | 1,4% ± 0,4% | n = 7 |
| $C^{1235}$ | mut | mut | mut | WT | mut | 0,24 ± 0,044 | 1,5 ± 0,6 | 93,4% ± 4,9% | 0,31 ± 0,05 | 1,3% ± 1,2% | n = 6 |
| $C^{1245}$ | mut | mut | WT | mut | mut | 0,09 ± 0,015 | 1,1 ± 0,3 | 97,4% ± 2,2% | 0,11 ± 0,03 | 0,5% ± 0,2% | n = 5 |
| $C^{12345}$ | mut | mut | mut | mut | mut | 0,04 ± 0,007 | 0,4 ± 0,3 | 89,5% ± 10,7% | 0,05 ± 0,01 | 0,5% ± 0,3% | n = 6 |

SU1 mut & SU3 mut = $\beta_2^{N303V}$ ; SU2 mut & SU4 mut = $\alpha_1^{N307V}$ ; SU5 mut = $\gamma_2^{H318V}$

**Supplementary Table 1**

| Parameter | Value | Unit |
|---|---|---|
| $k_{on}$ | 1,00E+05 | $M^{-1}.s^{-1}$ |
| $k_{off}$ | 100 | $s^{-1}$ |
| $\beta$ | 2000 | $s^{-1}$ |
| $\alpha$ | 500 | $s^{-1}$ |
| $\delta_f^+$ | 0.12 | $s^{-1}$ |
| $\delta_f^-$ | 0.08 | $s^{-1}$ |
| $\delta_s^+$ | 0.1 | $s^{-1}$ |
| $\delta_s^-$ | 0.01 | $s^{-1}$ |
| $\gamma_f^4$ | 3.5 | |
| $\varepsilon_f^4$ | 0.5 | |
| $\gamma_s^4$ | 1 | |
| $\varepsilon_s^4$ | 3 | |
| $\gamma_f^5$ | 3.5 | |
| $\varepsilon_f^5$ | 0.5 | |
| $\gamma_s^5$ | 1 | |
| $\varepsilon_s^5$ | 3 | |

Concerted model parameters

**Supplementary Table 2**

| Parameter | Model Ia | Model Ib | Model II | Model II-β | Model III | Unit |
|---|---|---|---|---|---|---|
| $k_{on}$ | 1.0 E+05 | 1.0 E+05 | 1.0 E+05 | 1.0 E+05 | 1.0 E+05 | $M^{-1}.s^{-1}$ |
| $k_{off}$ | 100 | 100 | 100 | 100 | 100 | $s^{-1}$ |
| $\beta$ | 2000 | 2000 | 2000 | 2000 | 2000 | $s^{-1}$ |
| $\alpha$ | 500 | 500 | 500 | 500 | 500 | $s^{-1}$ |
| $\delta^+$ | 0.035 | 0.035 | 0.24 | 0.24 | 0.28 | $s^{-1}$ |
| $\delta^-$ | 0.106 | 0.106 | 0.13 | 0.13 | 0.64 | $s^{-1}$ |
| $\delta_3^+$ | 0.042 | 0.042 | 0.05 | 0.05 | 8.0 E-04 | $s^{-1}$ |
| $\delta_3^-$ | 1.05 E-03 | 1.05 E-03 | 3.0 E-04 | 3.0 E-04 | 5.0 E-03 | $s^{-1}$ |
| $\gamma$ | - | - | - | - | 100 | |
| $\varepsilon$ | - | - | - | - | 10 | |
| $c_3^+$ | 4 | 10 | 10 | 3 | 3 | |
| $c_3^-$ | - | - | 1 | 2 | 2 | |
| $c_{34}^+$ | - | - | - | 4 | 4 | |
| $c_{34}^-$ | - | - | - | 4 | 3 | |
| $c_4^+$ | 10 | 100 | 30 | 30 | 20 | |
| $c_4^-$ | - | - | 3 | 3 | 1 | |
| $c_5^+$ | 10 | 100 | 30 | 30 | 60 | |
| $c_5^-$ | - | - | 3 | 3 | 1 | |

**Supplementary Table 3**
